# Advancing Knock-In Approaches for Robust Genome Editing in Zebrafish

**DOI:** 10.64898/2025.12.10.693484

**Authors:** Anjelica Rodriguez-Parks, Ella Grace Beezley, Steffani Manna, Isabella Silaban, Sarah I Almutawa, Siyang Cao, Hossam Ahmed, Megan Guyer, Sean Baker, Mark P Richards, Junsu Kang

**Affiliations:** Department of Cell and Regenerative Biology, School of Medicine and Public Health, University of Wisconsin-Madison, Madison, WI, 53705, USA; Department of Animal and Dairy Sciences, University of Wisconsin-Madison, Madison, WI 53706, USA; UW Carbone Cancer Center, School of Medicine and Public Health, University of Wisconsin-Madison, Madison, WI, 53705, USA

## Abstract

Precise genome editing remains a major challenge in functional genomics, particularly for generating knock-in (KI) alleles in model organisms. Here, we introduce the mini-golden system, a versatile Golden Gate-based subcloning platform that enables rapid assembly of donor constructs containing homology arms and a gene of interest. This system offers a library of middle entry vectors including diverse genes, enhancing the preparation of donor minicircles for KI applications. Using the mini-golden system, we efficiently generated a *foxd3^CreER^* KI zebrafish line, allowing conditional recombination in neural crest cells. To further improve genome editing precision, we developed a synthetic exon-based donor template strategy combined with fluorescence screening. Using this approach, we successfully engineered a targeted isoleucine-to-valine substitution (Ile-to-Val) in *hbaa1.2*, one of the two adult hemoglobin alpha genes in zebrafish. Importantly, despite the high sequence similarity between *hbaa1.2* and its paralog *hbaa1.1*, our strategy specifically edited *hbaa1.2*, demonstrating the effectiveness of the synthetic exon approach. This method minimized undesired recombination and significantly improved the identification of lines carrying the edited genome. Together, we provide a robust toolkit for efficient and precise genome engineering in zebrafish, with broad applicability to other model systems.

## Introduction

The advent of genome editing techniques, such as CRISPR/Cas9, has revolutionized genome editing approaches, allowing for targeted knock-out (KO) and knock-in (KI) applications (Hwang et al., 2013; Auer et al., 2014; Sander and Joung, 2014). KOs, such as mutating or deleting DNA fragments, are relatively easy, compared to KIs which require more intricate and precise steps (Hoshijima et al., 2019; Prykhozhij and Berman, 2024). KI strategies typically involve providing a donor template containing the gene of interest flanked by homology sequences at 5’ and 3’ regions of the target site. When this donor template is delivered into cells along with Cas9 and sgRNA, either homology-directed repair (HDR) or nonhomologous end joining (NHEJ) mechanism facilitates the integration of the gene of interest at the target site (Auer et al., 2014; Ata et al., 2016; Burg et al., 2018; Gutierrez-Triana et al., 2018; Prykhozhij et al., 2018; Wierson et al., 2020; Almeida et al., 2021; Levic et al., 2021; Mi and Andersson, 2023; Zhang et al., 2023; Prykhozhij and Berman, 2024; Oikemus et al., 2025). Despite advancements in KI methodologies, further refinement is needed to improve precision and efficiency, enhancing the impact of genome editing on biological research.

We and other groups demonstrated that minicircles can serve as a donor template enhancing the KI efficiency (Suzuki et al., 2016; Keating et al., 2024). However, creating a donor template containing both homology arms (HAs) is laborious and time-consuming. Generating a donor construct for KI generation requires multiple steps to add 5’ and 3’ HAs and supplementary sequences, such as a selection marker, depending on the complexity of the plasmids. Here, we developed the mini-golden system, a convenient, efficient subcloning platform and plasmid library designed to boost preparation of diverse donor templates. Utilizing this mini-golden system, we successfully generated the *foxd3^CreER^* KI line, which allows conditional recombination in neural crest cells (NCCs). The mini-golden system allows for one-step customization of donor templates, providing a robust and versatile tool for genome editing.

The genetic similarity between zebrafish and human positions zebrafish as an invaluable model for studying human diseases (Howe et al., 2013). Recent advancement of base and prime editing techniques enables the introduction of human disease-causing mutations into the zebrafish genome (Rosello et al., 2021; Petri et al., 2022; Richardson et al., 2023), significantly promoting the application of zebrafish for human disease modeling. Furthermore, iterative improvements in base editing, including optimization of the editing window, expansion of the PAM compatibilities, and introducing new DNA binding domains, have increased flexibility and broadened the range of targetable loci (Rosello et al., 2022; Zheng et al., 2023; Qin et al., 2024; Liu et al., 2025; Zhong et al., 2025). Despite these advances, the typical editing window for base editing spans several base pairs, creating the possibility of unintended edits when identical nucleotides occur within that window. Also, the current base and prime editing approaches rely on PCR-based genotyping to identify lines carrying the edited genome, which introduces multiple procedural steps and increases the risk of missing true positives. To improve screening efficiency, we incorporate a fluorescence selection marker, greatly advancing the identification of positive larvae through simple microscopy. Additionally, we employ a synthetic exon approach to avoid undesired recombination, further increasing efficiency. By integrating a synthetic exon strategy with a robust fluorescence-based screening method, we establish a highly effective approach for introducing amino acid (a.a.) changes in zebrafish models. Our strategy offers valuable resources and methodology to the genome editing toolbox for zebrafish and other animal model communities.

## Results

### mini-golden system for convenient and prompt subcloning

The mini-golden system utilizes Golden Gate assembly (Engler et al., 2008) to simultaneously and directionally combine 5’ HA, a gene of interest, and 3’ HA (**Fig. 1A and Supplementary Fig. S1A**). The mini-golden system consists of 4 components: 5’ and 3’ HAs, a middle entry vector (pMC-ME), and a destination vector (pMC-Dest). Our mini-golden system uses the BsaI restriction enzyme to create 4-base overhangs, thus all four components must not carry additional BsaI sites. 5’ and 3’ HAs are prepared by PCR from genomic DNA followed by gel extraction purification. Any internal BsaI sites in HAs must be mutated via overlap extension PCR or other available methods. For the destination vector, we modified the previously available minicircle generating vector^17^ by mutating existing BsaI sites and adding two outward facing BsaI sites in the multi-cloning site. We also constructed multiple middle entry vectors (pMC-MEs) encoding several fluorescence proteins (EGFP, mCherry, BFP, mNeongreen, mScarlet), membrane localized fluorescence proteins, CreER, or NTRv2 (Sharrock et al., 2022) (**Fig. 1B and Supplementary Table S1**). The detailed protocol for mini-golden system is described in S**upplementary Information**. The mini-golden system greatly facilitates quick and convenient subcloning to prepare a donor minicircle for KI strategy.

**Figure 1.**
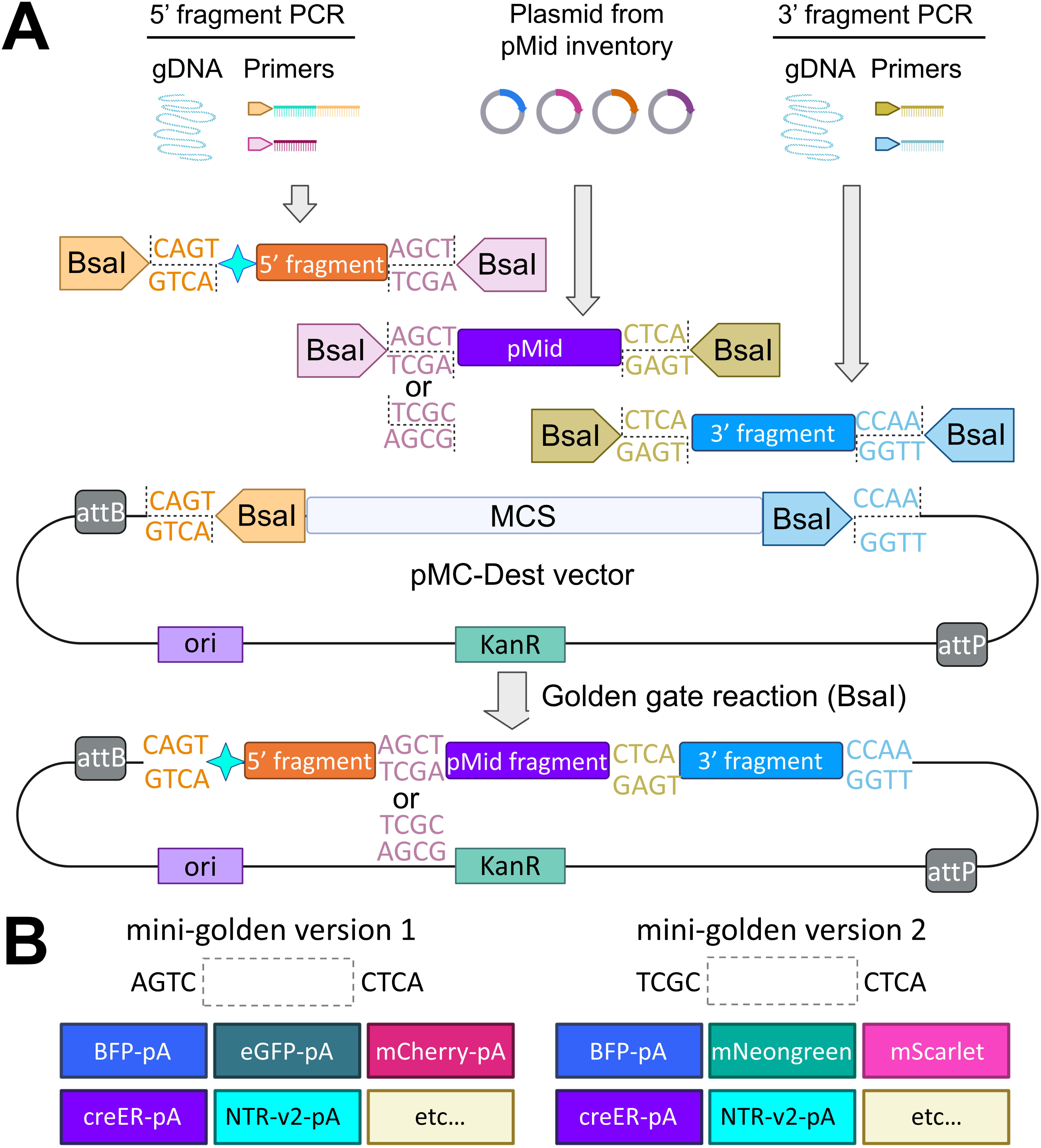
mini-golden system for robust creation of donor templates. (**A**) The schematic of the mini-golden system. The 5’ and 3’ homology arm (HA) fragments are prepared by PCR amplification followed by gel purification. Middle entry vector is chosen from the pMC-ME library. Linker sequences for mini-golden version 1 (Addgene Deposit 82577) and version 2 (Addgene Deposit 86145) are ACGT-CTCA and TCGC-CTCA, respectively. (**B**) pMC-ME middle entry vector library contains diverse constructs. The detailed protocol for mini-golden system is described in **Supplementary Information.**

### Generation of *foxd3^CreER^* knock-in strain via the mini-golden system

*Cre/loxp* system is a widely used genetic tool for spatiotemporal control of gene expression and lineage tracing(Sauer and Henderson, 1988; Liu et al., 2022). CreER, a fusion protein of Cre recombinase and the estrogen receptor (ER) ligand binding domain, enables temporal regulation through inducible activation. To generate a NCC *CreER* driver, we targeted *foxd3*, a pivotal transcription factor essential for NCC specification and differentiation (Lister et al., 2006; Stewart et al., 2006; Curran et al., 2009; Hochgreb-Hagele and Bronner, 2013; Candido-Ferreira et al., 2023), using our mini-golden system.

We designed our strategy to integrate *CreER* into the 5’ untranslated region (UTR) of *foxd3* and to delete a portion of the *foxd3* coding sequence to generate deletion mutants (**Fig. 2A and Supplementary Fig. S1B**). The sgRNA site is located approximately 830 bp upstream of the *foxd3* start codon. The 5’ HA fragment spans 826 bp starting from the sgRNA site, and the 3′ HA covered 547 bp beginning 403 bp downstream of the 5′ HA. From our mini-golden middle-entry library, we selected a plasmid containing *CreER* paired with an *FRT*-flanked *α-cry:mCherry* selection marker, which facilitates the identification of larvae carrying the modified genome. If necessary, *α-cry:mCherry* can be removed by introducing *flp* recombinase mRNA to prevent interference with *CreER* expression (Sugimoto et al., 2017). These components were assembled into a destination vector to create the donor plasmid. The final donor DNA was prepared using minicircle technology(Kay et al., 2010), which produces a minicircle DNA containing necessary DNAs without the bacterial backbone sequences (**Supplementary Fig. S1A**). Our previous work demonstrated that minicircle significantly enhances KI efficiency compared to a plasmid (Keating et al., 2024).

**Figure 2.**
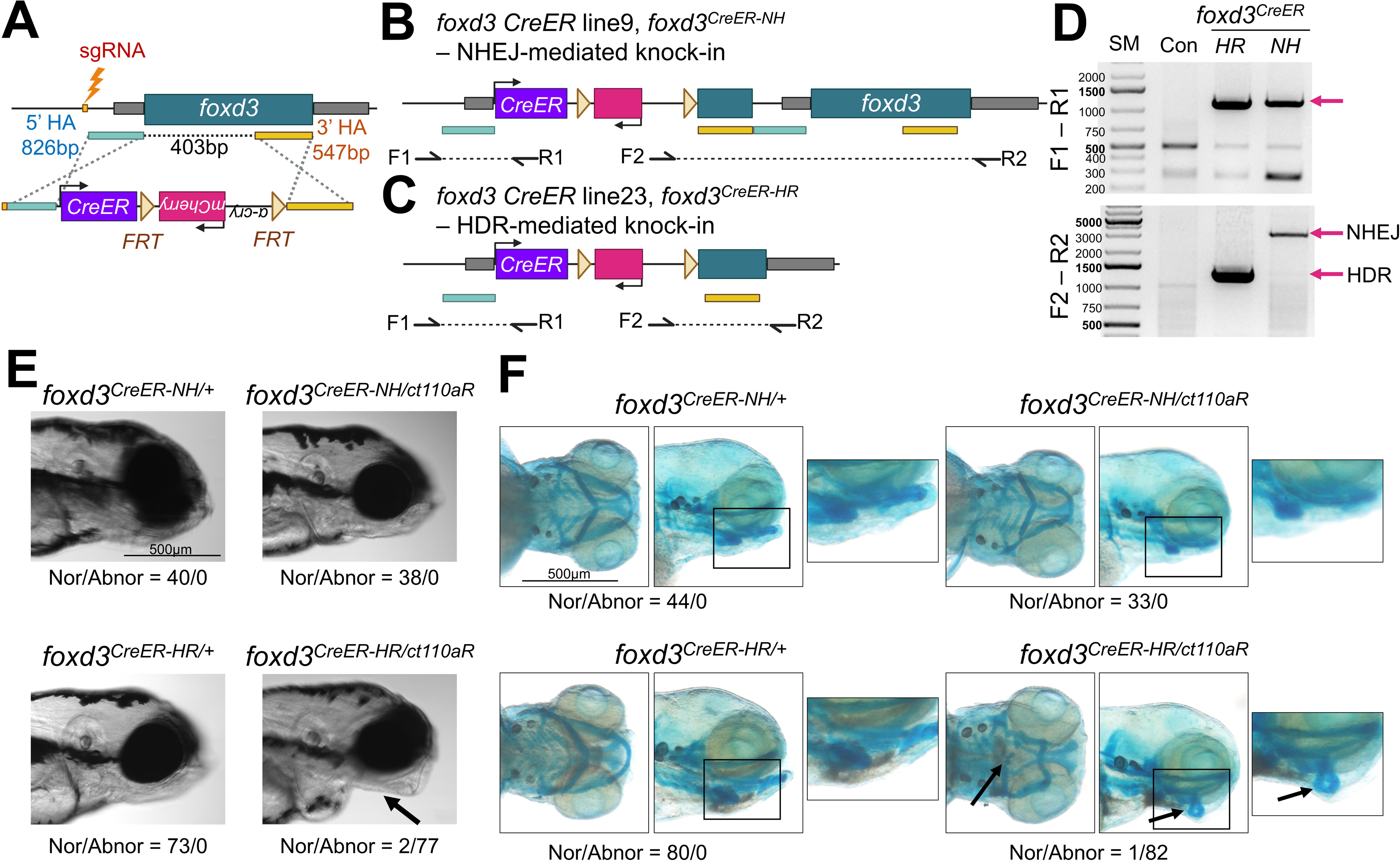
Generation of *foxd3CreER* knock-in (KI) line. (**A**) Schematic of genome editing strategy to create *foxd3^CreER^* KI line. (**B, C**) Schematic of *foxd3^CreER-NH^* (**B**) and *foxd3^CreER-HR^* (**C**). *foxd3^CreER-NH^* was created via non-homologous end joining (NHEJ)-mediated repair, resulting in an intact *foxd3* gene sequence. In contrast, *foxd3^CreER-HR^* was generated via homology-directed repair (HDR), leading to deletion of the 5′ portion of *foxd3*. (**D**) PCR analysis to genotype upstream (F1-R1) and downstream (F2-R2) regions of the integrated sites at the *foxd3* locus. (**E**) Lateral view of control heterozygotes and compound heterozygotes of the *foxd3^CreER-NH^*or *foxd3^CreER-HR^* and the *foxd3* enhancer trap line (*ct110aR*) at 4 days post-fertilization (dpf). While *foxd3^CreER-HR/+^*, *foxd3^CreER-NH/+^* and *foxd3^CreER-NH/ct110aR^* develop jaws normally, *foxd3^CreER-HR/ct110aR^* exhibits craniofacial defects (arrow). Nor, Normal. Abnor, Abnormal. (**F**) Ventral and lateral views of pharyngeal cartilages stained with Alcian blue at 5 dpf. Pharyngeal cartilages and arches are intact in *foxd3^CreER-HR/+^*, *foxd3^CreER-NH/+^* and *foxd3^CreER-NH/ct110aR^*, whereas *foxd3^CreER-HR/ct110aR^* exhibits disrupted cartilage phenotype (arrow). Scale bar, 500µm

We co-injected the donor minicircle, sgRNA, and Cas9 into one-cell stage embryos and assessed genome editing efficiency. The rate of *α-cry:mCherry^+^* F_0_ larvae was 60.37% (99/164). Among *α-cry:mCherry^+^* F_0_, 2 out of 16 individual embryos exhibited the correct 5’ and 3’ junction sizes (**Supplementary Fig. S1C**). *α-cry:mCherry^+^* F_0_ larvae were grown to adult, and we screened lens mCherry expressing fish again at the adult stage. From ∼30 *α-cry:mCherry^+^* F_0_ adult, we established 7 lines transmitting *α-cry:mCherry^+^* to the next generation (**Fig. 2B-D and Supplementary Fig. S1D**). PCR analysis of F_1_ animals revealed that 4 lines failed to amplify both 5’ and 3’ flanking regions. Line 8 showed the expected 3’ fragment but a longer 5’ fragment, while line 9 produced the expected 5’ fragment but a longer 3’ fragment (**Fig. 2B, D and Supplementary Fig. S1D**). Line 23 generated correctly sized bands at both junctions (**Fig. 2C, D**). Sanger sequencing analyses confirmed that *CreER* was integrated into the *foxd3* locus via NHEJ in line 9, hereafter referred to as *foxd3^CreER-NH^*, (**Fig. 2B, D**). By contrast, line 23 exhibited homology-directed repair (HDR)-mediated integration, hereafter referred to as *foxd3^CreER-HR^*, (**Fig. 2C, D**).

*foxd3* is an important transcription factor for pre-migratory NCC specification and fate determination, and *foxd3* mutants display deformed jaws (Lister et al., 2006; Stewart et al., 2006; Hochgreb-Hagele and Bronner, 2013). To assess whether *CreER* integration affects *foxd3* function, we crossed *foxd3^CreER-NH^* or *foxd3^CreER-HR^* with G*t(foxd3:mcherry)^ct110aR^*, a *foxd3* gene trap reporter and loss-of-function allele (Hochgreb-Hagele and Bronner, 2013). While *foxd3^CreER-NH/ct110aR^* exhibited no craniofacial abnormalities, we observed that *foxd3^CreER-HR/ct110aR^* displayed jaw malformations (**Fig. 2E**). All heterozygote controls have normal shaped jaws. Alcian Blue staining revealed the defects in the posterior pharyngeal elements of *foxd3^CreER-HR/ct110aR^*, but not in *foxd3^CreER-NH/ct110aR^* and heterozygote controls (**Fig. 2F**). Moreover, Reverse Transcription Polymerase Chain Reaction (RT-PCR) analysis confirmed that functional *foxd3* transcripts were undetectable in cDNA extracted from *foxd3^CreER-HR/ct110aR^*, whereas *foxd3^CreER-NH/ct110aR^* has it (**Supplementary Fig. S2**). Collectively, our mini-golden-mediated strategy successfully generated a *CreER* KI driver for *foxd3. foxd3^CreER-HR^* represents a complete *foxd3* loss-of-function allele, while *foxd3^CreER-NH^* retains the endogenous *foxd3* activity. These strains provide valuable tools for NCC lineage studies in zebrafish.

### *foxd3^CreER^* and tamoxifen-mediated lineage tracing strategy labels neural crest-derived cells

To determine *foxd3* expression pattern, we analyzed the G*t(foxd3:mcherry)^ct110aR^* gene trap line (Hochgreb-Hagele and Bronner, 2013). Consistent with previous reports (Hochgreb-Hagele and Bronner, 2013), our confocal imaging confirmed mCherry expression in cranial NCCs and peripheral glial cells (**Supplementary Fig. S3A, B**). Additionally, as shown in previous studies, we observed mCherry expression in the paraxial mesoderm starting at 2 days post-fertilization (dpf), which declined by 5 dpf (**Supplementary Fig. S3A, B**) (Odenthal and Nusslein-Volhard, 1998; Lee et al., 2006; Stewart et al., 2006). While the *α-cry:mCherry* selection marker drives lens mCherry expression in both *foxd3^CreER-NH^* and *foxd3^CreER-HR^*, *foxd3^CreER-HR^* exhibits additional mCherry expression in the paraxial mesoderm (**Supplementary Fig. S3C, D**). This unexpected expression pattern of the selection marker indicates the presence of the potential mesoderm regulatory elements downstream of *foxd3*, which may influence *α-cry:mCherry* transcription for the *foxd3^CreER-HR^*. *flp* recombinase mRNA injection into the one-cell stage embryos successfully delete the *FRT*-franked *α-cry:mCherry* cassette, resulting in the lack of lens expression (**Fig. 3 and S3E, F**).

**Figure 3.**
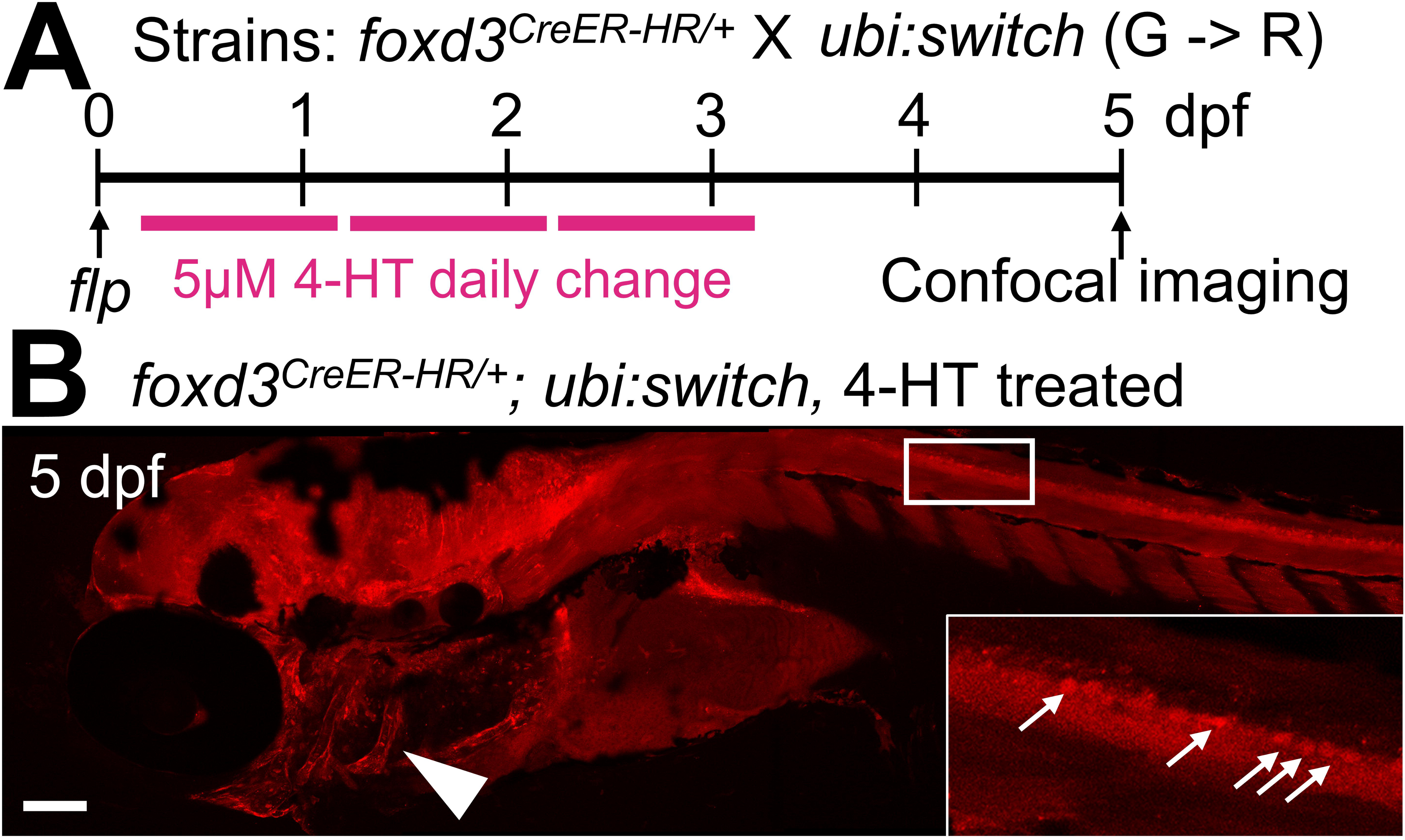
*foxd3^CreER-HR^* KI induces recombination in neural crest cells (NCCs) **(A)** Schematic of 4-hydroxytamoxifen (4-HT) treatment with *foxd3^CreER-HR/+^; ubi:switch* (Green to Red). *flp* mRNA was injected to remove the FRT-franked *α-cry:mCherry* cassette. (**B**) Representative confocal images of zebrafish larvae at 5 dpf following 4-HT treatment. The *ubi:switch* transgene undergoes recombination in NCCs upon 4-HT-induced activation of *CreER* in *foxd3*-expressing NCCs, labeling craniofacial NCC-derived cells (arrowhead) and peripheral glial cells (arrow). Insets show magnified views of peripheral glial cells in 4-HT-treated larvae (**B**). Scale bar, 100µm

To validate whether newly generated *foxd3^CreER-HR^* can drive expression of functional *CreER* in NCCs, we crossed *foxd3^CreER-HR^*with *ubi:switch* (Mosimann et al., 2011) and examined whether NCCs were specifically labelled following tamoxifen treatment. *flp* mRNA was injected at the one-cell stage embryos to prevent *CreER*-mediated mCherry expression. 4-HT was treated from 5 to 77 hours post-fertilization (hpf) to induce recombination and mCherry expression was evaluated to determine *CreER* expressing cells. At 5 dpf, mCherry expression was detectable in cranial NCCs and peripheral glial cells, matching to that of G*t(foxd3:mcherry)^ct110aR^* (**Fig. 3 and Supplementary Fig. S3A, B**). These data indicate successful *CreER*-mediated-recombination of *foxd3^CreER-HR^* in NCCs to label NCC-derived cells.

### Adult hemoglobin alpha chain gene structures in zebrafish

Hemoglobin (Hb), a key oxygen-transporting protein in red blood cells, is composed of two alpha- and two beta-globin chains (Chan et al., 1997; Brownlie et al., 2003). While Hbs across animal species share a similar structural architecture, variations in certain a.a. residues lead to differences in oxygen-carrying capacity. For instance, fetal Hbs exhibit a higher affinity for oxygen than adult Hbs (Manning et al., 2020). In most fish Hbs, an isoleucine (Ile) residue occupies position 11 of the E-helix (E11), whereas mammalian Hbs have smaller amino acids at this position, such as valine (Val) (**Supplementary Fig. S4A**). Previous studies suggest that IleE11 may promote the dissociation of the neutral superoxide radical, potentially enhancing Hb oxidation (Aranda et al., 2009). To investigate this, we aim to substitute IleE11 to ValE11 in zebrafish.

Zebrafish genome encodes two adult alpha-globin genes(Brownlie et al., 2003), including *hbaa1* and *si:ch211-5k11.8,* which we have annotated as *hbaa1.1* and *hbaa1.2*, respectively (**Fig. 4A**). Both genes carry Ile at the E11 (**Supplementary Fig. S4A, B**). To determine the major adult isoform, we collected blood from adult fish and performed mass spectrometry. Although peptides from both Hbaa1.1 and Hbaa1.2 were detected, Hbaa1.2 showed slightly higher expression levels (**Supplementary Fig. S4C, D**). Based on this finding, we selected Hbaa1.2 for modification of the IleE11 to ValE11 residue. *hbaa1.1* and *hbaa1.2* exhibit extremely high sequence identity at the a.a., coding, and genomic DNA levels (**Fig. 4B**). Although most sgRNA sites target both genes, we identified an efficient sgRNA site specific to *hbaa1.2,* which is positioned 63 base pairs upstream of ATC (I64) codon. This site exhibits two base pair differences compared to the corresponding site of *hbaa1.1* (**Fig. 4C, D**).

**Figure 4.**
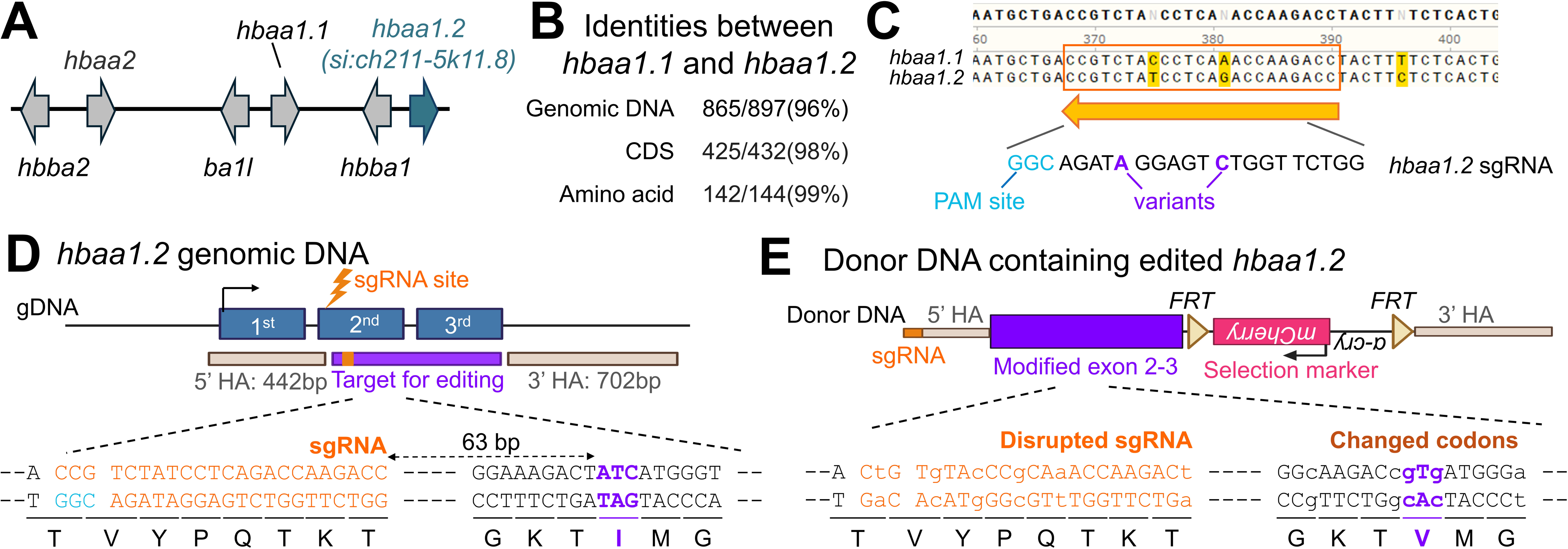
Design of a single amino acid (a.a.) substitution in Hbaa1.2. (**A**) Schematic diagram of the α- and β-globin gene clusters in zebrafish. (**B**) Sequence similarity between *hbaa1.1* and *hbaa1.2* at genomic DNA, coding sequence (CDS), and a.a. level. (**C**) sgRNA target site in *hbaa1.2* and the homologous sequence in *hbaa1.1*. Nucleotide variants between these two genes are highlighted by red. (**D**) *hbaa1.2* gene structure, sgRNA sequence (orange), and the codon encoding I residue targeted for the substitution (ATC, red). Corresponding a.a. sequences are shown below each codon. (**E**) Structure of the donor minicircle and the synthetic exon sequence designed for *hbaa1.2* editing. Individual codons, except the one encoding the target valine (V) residue, were replaced with synonymous codons to maintain protein sequence while preventing re-cutting by Cas9 as well as undesired recombination.

### Synthetic exon-based genome editing combined with fluorescence screening enables a single amino acid substitution in Hbaa1.2

A primary challenge in genome editing for a.a. substitution is the low efficiency of selecting animals that carry the desired modification. Previous a.a. editing approaches, including single oligo-mediated and prime/base editing, relied on PCR-based screening to identify animals carrying the edited genome (Rosello et al., 2021; Petri et al., 2022; Rosello et al., 2022; Richardson et al., 2023). Given the low efficiency of genome editing techniques and a reduced germline transmission rate, this PCR-based approach may fail to select positive lines due to the complicated multiple steps for genotyping and dilution of the edited genome by massive amount of unedited wild-type genomic DNA. Additionally, this process is labor-intensive and time-consuming. To address these limitations, we incorporated a fluorescence selection marker to simplify screening process. This approach allows for the identification of potential positive lines using fluorescence microscopy (**Fig. 5A**). We used *FRT*-flanked *α-cry:mCherry,* which drives lens expression, as a selection marker (**Fig. 4E**). If a selection marker interferes with *hbaa1.2* gene expression, it can be eliminated by introducing the *flp* recombinase mRNA.

**Figure 5.**
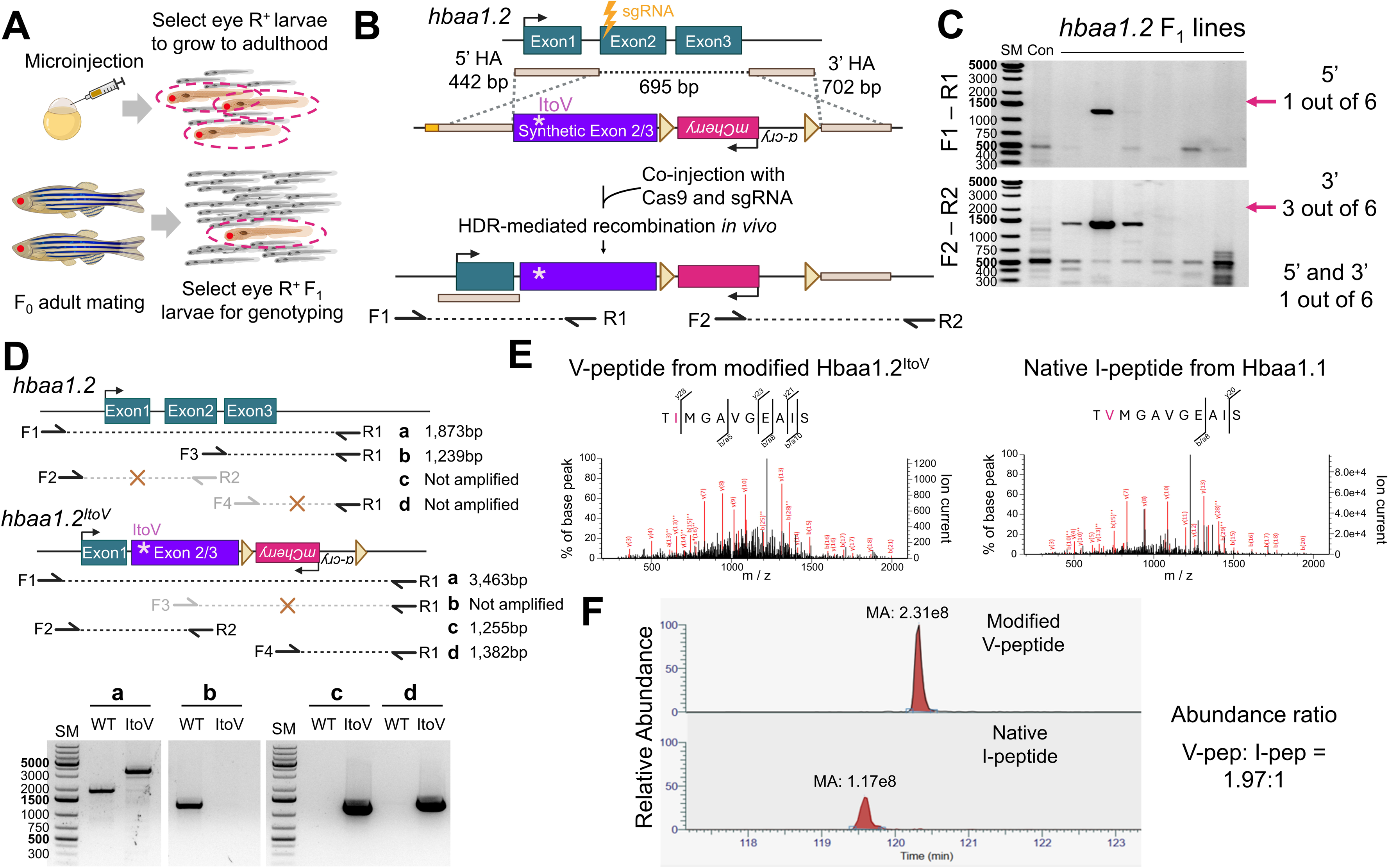
Synthetic exon-mediated genome editing combined with fluorescence-based screening successfully engineers the single amino acid of Hbaa1.2. (**A**) Fluorescence-based screening enriches for larvae carrying the edited genome, facilitating efficient identification of correctly modified individuals. (**B**) Schematic of genome editing strategy to substitute I with V in Hbaa1.2. (**C**) PCR genotyping of F_1_ stable lines targeting the upstream (F1–R1) and downstream (F2–R2) regions flanking the integration site at the *hbaa1.2* locus. (**D**) PCR analysis comparing wild-type (WT) siblings and *hbaa1.2^ItoV^* homozygotes (ItoV) to distinguish wild-type and *hbaa1.2^ItoV^* alleles. (**E**) Mass spectrometry analysis of blood from *hbaa1.2^ItoV^* homozygote fish. The edited peptide containing I to V substitution (V-pep) is detectable. (**F**) Relative abundance of the edited V-pep versus the native I-pep.

The target a.a. is located within the 2^nd^ exon (**Fig. 4D**) and a selection marker must be placed in a non-coding region to avoid disrupting *hbaa1.2* expression. To address these challenges, we positioned the selection marker downstream of *hbaa1.2* (**Fig. 5B and Supplementary Fig. S5A**). However, this configuration can lead to undesired recombination events between one of HA and DNA sequences surrounding the target a.a.. To prevent unwanted recombination, we devised the use of a synthetic exon as a donor template. The sgRNA site and downstream coding DNA, except I64V in the donor, are altered with synonymous substitution to minimize undesired recombination (**Fig. 4E and Supplementary Fig. S5A)**. However, the 3’ UTR region was left unmodified to retain potential regulatory elements. Taking these considerations, our donor template contains the following elements in order: a sgRNA site, a 442 bp 5’ HA, a modified partial 329 bp *hbaa1.2* CDS having I64V change, 279 bp 3’ UTR and pA, a FRT-flanked lens-specific selection marker, and a 702 bp 3’ HA (**Fig. 4E and 5B**). Using this synthetic exon-based approach, we aim to precisely engineer the IleE11 to ValE11 substitution in the Hbaa1.2, enhancing the efficiency of positive line identification and successful genome modification.

We co-injected the donor minicircle, sgRNA, and Cas9 and evaluated genome editing with F_0_. The rate of *α-cry:mCherry^+^*F_0_ larvae is 45.31% (29/64). We sorted *α-cry:mCherry*^+^ larvae and performed genotyping to assess recombination events. Of these, 77% *α-cry:mCherry*^+^ larvae (17/22) showed amplification of the 3’ junction, but 22% (5/22) had successful integration of the modified *hbaa1.2* coding DNA at both 5’ and 3’ junctions (**Supplementary Fig. S5B**). The higher rate of 3’-only recombinants suggests unintended recombination between the 285 bp 3’ UTR and 3’ HA, rather than 5’ and 3’ HA. We raised *α-cry:mCherry*^+^ F_0_ embryos to adulthood and screened their progeny for *α-cry:mCherry* expression. From approximately 40 F_0_ individuals, we identified six founders. Of these, two exhibited incomplete KI, likely due to recombination between the 3′ UTR-pA and the 3′ HA, resulting in failure to integrate the 5′ synthetic exon (**Fig. 5C**). One founder carried the fully and accurately edited genome with correct integration at *hbaa1.2* locus (**Fig. 5C, D**). PCR analysis with primers specific to wild-type and edited *hbaa1.2* (*hbaa1.2^ItoV^*) revealed that wild-type primers amplified *hbaa1.2* fragment from genomic DNA extracted of wild-type sibling but failed to amplify the fragment from *hbaa1.2^ItoV^* homozygotes. Conversely, *hbaa1.2^ItoV^* specific primers amplified the target fragment only in the *hbaa1.2^ItoV^* homozygotes (**Fig. 5D**). These results demonstrate that our strategy can achieve precise genome editing even in the presence of highly similar endogenous sequences.

To determine whether the edited *hbaa1.2^ItoV^* allele produces a protein without disrupting expression, we collected blood from adult *hbaa1.2^ItoV^* homozygous fish and performed mass spectrometry analysis. We successfully detected the peptides carrying the I-to-V substitution (**Fig. 5E**). Because *hbaa1.1* remains unmodified in *hbaa1.2^ItoV^* homozygotes, native I-containing peptides were also detectable. Notably, quantitative analysis revealed a higher abundance of V-containing peptides compared to I-containing ones, indicating that the edited *hbaa1.2^ItoV^* allele is actively expressed (**Fig. 5F**). These results suggest that our strategy for single a.a. substitution can serve as an effective genome editing approach.

## Discussion

CRISPR/Cas9-mediated KIs rely on inducing a targeted DNA double-strand break (DSB) and providing a donor template for integration. Assembling donor templates - by combining two HAs with a gene of interest into a single plasmid - requires heavy subcloning. To simplify labor-intensive cloning steps, we developed a mini-golden-mediated subcloning strategy, which enables rapid and efficient construction of donor templates. A diverse set of middle entry vectors supports the construction of donor templates carrying various genes of interest, including fluorescence protein with subcellular organelle targeting sequences, the *CreER* recombinase, the *NTR* gene for genetic ablation, and more. In addition, Golden Gate subcloning allows the flexible incorporation of auxiliary DNA modules between the middle entry fragment and either the 5′ or 3′ HA, offering enhanced flexibility in construct design. Middle entry fragments can also be assembled from multiple modular components - ideal for generating fusion proteins - thus expanding the versatility of this system. Our strategy requires to mutate BsaI sites within HAs for the Golden Gate assembly. Although the impact of sequence mismatches on HDR-mediated KI is not fully understood, we successfully created the *foxd3^CreER^* KI line despite the presence of a BsaI mutation in the 5’ HA. Together with recent studies showing that short HA (40-50bp) can support efficient HDR-mediated KI (Wierson et al., 2020; Mi and Andersson, 2023; Oikemus et al., 2025), our results suggest that HDR can tolerate limited sequence mismatches within the longer HAs without substantially compromising KI efficiency. To ensure compatibility with the established KI methods such as GeneWeld (Wierson et al., 2020), we also provide destination vectors carrying a universal sgRNA site at the 5′ end. Furthermore, the middle entry library can serve as a template for generating short-arm donor fragments containing 5′ modifications, such as C6 linker (AmC6) (Mi and Andersson, 2023), 2’O-methyl-end modification (2’OMe) (Oikemus et al., 2025), biotin (Gutierrez-Triana et al., 2018; Zhang et al., 2023) or other chemical modification. Altogether, we present the mini-golden system as a flexible and efficient toolkit for CRISPR-based genome editing applications.

Early strategies for introducing small edits, such as single nucleotide variant or short DNA insertion, utilized single-stranded oligodeoxynucleotides (ssODNs) containing both HAs and variant sequence (Burg et al., 2018; Prykhozhij et al., 2018; Tessadori et al., 2018). More recent advances in genome editing include prime and base editing approaches. Prime editing employs a Cas9 nickase fused to a reverse transcriptase and a prime editing guide RNA (pegRNA) to directly write new sequences into the genome without donor templates (Petri et al., 2022). Similarly, base editing (using cytidine or adenine deaminase fused to Cas9) has been applied in zebrafish to precisely convert single bases without inducing DSB (Rosello et al., 2021). In particular, the recent advancement of base editing techniques has robustly improved efficiency (Zheng et al., 2023; Qin et al., 2024; Liu et al., 2025; Oikemus et al., 2025; Zhong et al., 2025). However, these approaches still have some limitations, including unintended conversion of bystander nucleotides and a limited set of validated conversion options. To date, C-to-T and A-to-G edits have been shown to be effective in zebrafish, whereas efficient conversion of other types has not yet been confirmed (Liu et al., 2025). Another challenge is how to effectively identify animals carrying the edited genome. Our strategies address this challenge in two important ways. First, our strategy enables convenient and robust enrichment of larvae harboring edited genome using fluorescence signal. Our method allows reliable identification of edited animals simply by monitoring fluorescent reporter expressions, thereby focusing downstream analysis on animals with the edits. This also improves genotyping sensitivity by avoiding dilution of edited gDNA with that of unedited animals.

Second, our synthetic exon approach minimizes undesired recombination events. Previous studies to insert two *loxp* sites also used fluorescent screening enrichment (Shin et al., 2023). However, those approaches placed wild-type sequences between two HAs, resulting in low efficiency due to unwanted recombination between either HA and the homologous endogenous target exon. Our synthetic exon-mediated strategy can mitigate this by avoiding direct sequence homology with the endogenous exon. Nevertheless, codon replacement within the synthetic exon may lead to unexpected outcomes, such as affecting mRNA stability (Wu and Bazzini, 2023). To minimize such effect, optimal synonymous codon substitutions should preserve the native codon usage balance. To support this, we provide practical guidelines and templates (**Fig. 6 and Supplementary Data**). While our edited *hbaa1.2* allele is expressed, further validation with other gene targets will be necessary to fully optimize the synthetic exon-mediated genome editing approach.

**Figure 6.**
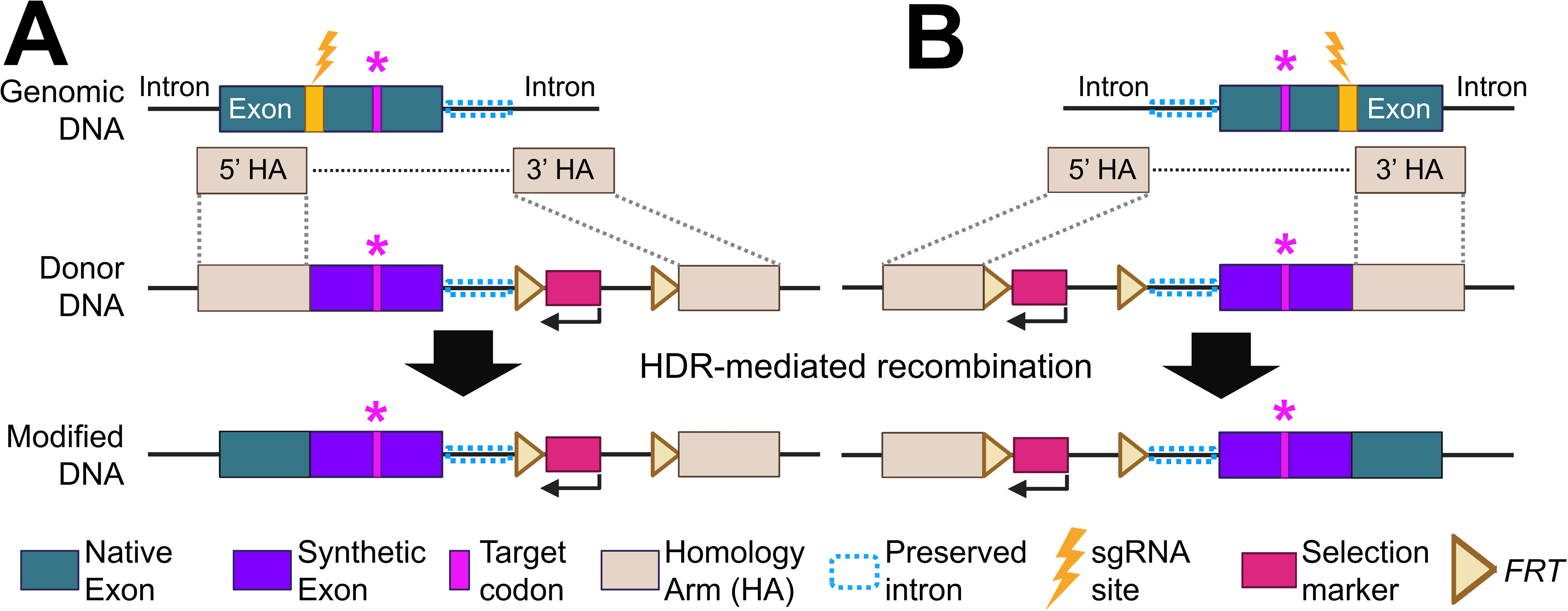
The schematic of the proposed design for single amino acid substitution. A recent study reported that placing HAs more than 5 bp from the cut site reduces recombination efficiency (Oikemus et al., 2025). Our data demonstrate that recombination can occur efficiently even when two HAs are positioned distantly. Taken together, these findings suggest that at least one HA may need to be placed within 5 bp of the cut site to maintain high efficiency. Based on this framework, we propose two potential design strategies. (**A**) If the sgRNA site is located upstream of the target codon, the 5’ HA ends at the cut site and the *FRT*-franked fluorescence selection marker can be inserted within the following intron in the reverse-complement orientation. (**B**) If the sgRNA site is located downstream of the target codon, the 3’ HA begins at the cut site and the *FRT*-franked fluorescence selection marker can be inserted into the preceding intron in the reverse-complement orientation. In both cases (**A**, **B**), codons between the sgRNA cut site and the end of exon, excluding the target codon, will be replaced with synonymous substitutions. Because exon-intron junctions contain key regulatory elements, we recommend preserving 30-40 bp of the native intronic sequence to avoid disrupting splicing.

Overall, our strategy offers an efficient method to generate KI lines, expanding the capacity to create KI animals and enabling deeper investigation into gene function across diverse biological processes.

## Materials and Methods

### Zebrafish

Wild-type or transgenic male and female zebrafish of the outbred Ekkwill (EK) or AB strain ranging up to 18 months of age were used for all zebrafish experiments. The following transgenic lines were used in this study: *Gt(foxd3:mCherry)^ct110aR^ (Hochgreb-Hagele and Bronner, 2013), Tg(–3.5ubi:loxP-EGFP-loxP-mCherry)^cz1701Tg^ or ubi:switch (Mosimann et al., 2011), foxd3^CreER#9^ (uwk45), foxd3^CreER#23^ (uwk46),* and *hbaa1.2^ItoV^(uwk47).* The water temperature for adult animals was maintained at 26°C unless otherwise indicated. Embryos and larvae were maintained at 28°C in egg water containing 300 mg/L sea salt, 75 mg/L calcium sulfate, 37.5 mg/L sodium bicarbonate, and 0.0001% methylene blue. Animals were anesthetized in 0.02% tricaine until gill movement stopped. *flp* mRNA was generated using the pCS2-Flp plasmid and the mMessenger mMachine SP6 kit (ThermoFisher Scientific). For 4-hydroxytamoxifen (4-HT) treatments, zebrafish larvae were incubated with 5 uM of 4-HT (Sigma-Aldrich, H7904) for 3 consecutive days from 5 to 77 hpf with daily media replacement. Work with zebrafish was performed in accordance with University of Wisconsin-Madison guidelines.

RNA was isolated from larvae using Tri-Reagent (ThermoFisher). Complementary DNA (cDNA) was synthesized from 300 ng to 1 μg of total RNA using a NEB ProtoScript II first strand cDNA synthesis kit (NEB, E6560). Primer sequences used for RT-PCR are listed in **Supplementary Data File 1**.

### Subcloning and transgenesis

mini-golden destination vector: To generate the pMC-Dest-BsaI destination vector (pMC-Dest-BsaI; Addgene ID: 200550), three BsaI sites in the pMC.BESPX-MCS2 (MN100B-1, SBI) were mutated using a site-directed mutagenesis kit (NEB, E0554S). A double-stranded oligonucleotide that contains two BsaI sites producing CTCA and CCAA overhang at the 5’ and 3’, respectively, was inserted via EcoRI and SalI enzymes-mediated subcloning. A recent study reported that addition of a reporter cassette outside of the HAs can aid in detecting off-target insertions (Shin et al., 2023). To incorporate this strategy, we further modified the destination vector by inserting a *cmlc2:mCherry-pA* cassette via PspOMI and SalI enzymes-mediated subcloning (pMC-Dest-BsaI-cmlc2:mCherry; Addgene ID: 241213). To expand the mini-golden system for compatibility with the GeneWeld method(Wierson et al., 2020), we also added a universal sgRNA sequence (GGGAG GCGTT CGGGC CACAG CGG) at the 5’ end of the cloning site via PspOMI and SalI enzymes-mediated subcloning (pMC-Dest-uni-gRNA5-BsaI; Addgene ID: 241150). Primers used for subcloning are listed in **Supplementary Data File 1**.

Middle entry library vector version 1: The middle entry vector backbone was derived from Addgene plasmid #64247 (pU6a:sgRNA#3) (Yin et al., 2015), which carries a spectinomycin resistance gene. pU6b:sgRNA#3 was a gift from Wenbiao Chen (Addgene plasmid #64247; http://n2t.net/addgene:64247; RRID:Addgene_64247). Two new BsaI sites generating AGTC and CCAA overhangs at the 5’ and 3’ end, respectively, replaced the *pU6a* promoter sequence via PspOMI and XbaI enzymes-mediated subcloning. Insert fragments were prepared by PCR with primers tagged with BsaI site to produce AGTC and CCAA overhang at the 5’ and 3’ end, respectively. After digestion of the vector and the insert with BsaI, linearized fragments were purified using a gel extraction kit (Takara Bio, 740609.250), ligated with T4 ligase (NEB, M0202M), and transformed into *E.coli* for further subcloning. An “AGTC” linker sequence remains between the 5’ HA and a gene of interest.

Middle entry library vector version 2: We are aware that “AGTC” could potentially act as a splicing acceptor (AG) or donor (GT) site, possibly leading to unintended splicing events when integrated into the genome. To address this concern, we resubcloned middle entry vectors using TCGC and CTCA overhangs at 5’ and 3’ end, respectively. Plasmids are listed in **Supplementary Table S1.** Primers used for subcloning are listed in **Supplementary Data File 1**.

*foxd3 CreER* donor construct: 5’ and 3’ HAs were amplified using genomic DNA extracted from fish that were used for injection. The corresponding sgRNA site for *foxd3* was positioned at the 5’ end of the 5’ HA. The sgRNA and PAM sequences for *foxd3* are “GCACA GGTGA GCGAC GCATG TGG”. These HA fragments and pMC-ME_*CreER-pA-FRT-acry:mCherry-pA-FRT* were assembled with pDest-BsaI via BsaI-mediated golden gate reaction. To purify the minicircle vectors, parental plasmids were transformed into the ZYCY10P3S2T *E. coli* Minicircle producer strain (MN900A-1, SBI). Minicircles were prepared as described in the (Keating et al., 2024). Primers used for subcloning are listed in **Supplementary Data File 1**.

*hbaa1.2* modified donor construct: The sgRNA and PAM sequences for *hbaa1.2* are “GGTCT TGGTC TGAGG ATAGA CGG”. The fragment containing sgRNA, 5’ HA, and modified exon sequence was synthesized as gBlocks by IDT and subcloned into pMC.BESPX-MCS2 via EcoRI and ClaI enzymes-mediated subcloning. The 3′ HA was subsequently amplified from genomic DNA extracted from the fish used for injection and inserted into the vector via BamHI and SacI enzyme-mediated subcloning. Finally, the FRT-flanked *α-cry:mCherry-pA* fragment was inserted in the reverse complement orientation using BamHI and ClaI enzyme-mediated subcloning. To purify the minicircle vectors, parental plasmids were transformed into the ZYCY10P3S2T *E. coli* Minicircle producer strain (MN900A-1, SBI). Minicircles were prepared as described in (Keating et al., 2024). Primers used for subcloning are listed in **Supplementary Data File 1**.

### Generation of *foxd3^CreER^* and *hbaa1.2^ItoV^* KI lines and genotyping

sgRNAs were synthesized by a cloning-free method as described in (Varshney et al., 2015). Briefly, primers containing T7 promoter, sgRNA target and partial sgRNA scaffold sequences were annealed with the universal sgRNA 3’ scaffold primer, and the annealed oligos were filled using the thermocycler and PCRBio polymerase (Genesee Scientific) to generate sgRNA templates. sgRNAs were synthesized using the HiScribe T7 kit (NEB, E2050S) and purified by the RNA purification Kit (Zymogen, R1016) according to the manufacturer’s instructions. A sgRNA (25∼30 ng/ul) and a donor minicircle (20-25 ng/ul) were mixed and co-injected with Cas9 protein (0.5ug/ul; PNABio, CP01) into the one-cell stage embryos. Genomic DNA was extracted, and KI was confirmed by genotyping. Primers used for genotyping are listed in **Supplementary Data File 1**. Larvae having mCherry in lens were sorted and raised to adulthood, and founders were screened with F_1_ progenies. The founder was outcrossed with wild-type animals to generate heterozygous animals.

### Alcian Blue Cartilage Staining

To assess cartilage development, we used an acid-free Alcian blue staining method(Balasubramanian et al., 2023) on 5 dpf larvae. Larvae were first washed in PBS and then euthanized by cold-shock on ice for 10 minutes. Larvae were transferred to 1.5 mL microcentrifuge tubes containing 4% paraformaldehyde (PFA; VWR, 102091-904) and incubated for two nights at 4 °C. Following fixation, samples were rinsed twice in PBST (PBS + 0.1% Tween-20) for 5 minutes each. Larvae were dehydrated in 50% ethanol and rocked at room temperature for 10 minutes. Ethanol was replaced with 1mL of Alcian blue staining solution (final concentrations of: 0.1mg/mL Alcian blue in methanol, 60mM MgCl2, and 70% ethanol). Samples were rocked overnight at room temperature and protected from light using foil. Stained larvae were rinsed once in PBST then transferred to a 24-well culture plate containing 2 mL bleaching solution (final concentrations of: 3% H_2_O_2_ and 0.5% KOH). Samples were incubated for 30 minutes in the dark. Following bleaching, larvae were rinsed again in PBST. Larvae were cleared by rocking in 25% glycerol with 0.25% KOH for 1 hour, followed by incubation overnight in 50% glycerol with 0.25% KOH. Larvae were stored in 50% glycerol with 0.1% KOH at 4°C.

### Imaging

Whole-mount larval images were acquired using an AxioZoom stereo fluorescence microscope (Zeiss) or an Olympus FV3000 confocal microscope (Olympus). Whole-mount imaging for Alcian blue stained larvae was acquired using an AxioZoom and color camera. Further image processing was carried out manually using Zen (Zeiss), Fluoview (Olympus), Photoshop, or FIJI/ImageJ software.

### Blood collection and Mass spectrometry

Our blood collection method was adapted from (Babaei et al., 2013). A 22G needle was used to bore a single hole in the bottom of a 0.6 mL centrifuge tube, followed by four additional equidistant holes made with a 27G needle around the central hole. This modified 0.6 mL tube was then placed into a 1.5 mL centrifuge tube. A 15 µL drop of 20 U/mL heparin in PBS was added to the side of the inner 0.6 mL tube. Fish weighing more than 0.4 g were anesthetized with 0.02% tricaine (MS222), then gently dried with a paper towel to remove excess water. Using a straight blade dipped in heparin solution, the fish tails were amputated just above the base of the caudal fin. The fish were quickly transferred cut-side down, into the 0.6mL tube, ensuring that the cut tail contacted the heparin droplet. After closing the lid of the inner tube, samples were centrifuged at 60 x g for 5 minutes at 14°C. The collected blood was gently flicked to mix with the heparin solution to minimize clotting. Samples were kept on ice until used in downstream applications. Fish carcasses were subsequently used for genomic DNA (gDNA) and RNA extraction.

Hb peptides were prepared using trypsin/LysC and iodoacetic acid (Kassa et al., 2021). Dithiothreitol final concentration was 2 mM. Samples were desalted with a C18 OMIX tip. Samples were injected into a Pepmap C18, 3 μM, 100A, 75μM ID, 15cm reversed phase column. Samples were analyzed on an Orbitrap Elite coupled to an EASY-Spray ion source in data dependent MS/MS mode.

## Acknowledgments

We thank the UW–Madison SMPH BRMS staff and members of the Kang Lab for zebrafish care; Sarah C Kucenas and Marianne Bronner for *Gt(foxd3:mCherry)^ct110aR^*; Len Zon and Jingli Cao for *ubi:switch*; Rachel Wong and Owen Lawrence for sharing a membrane localization sequence containing plasmid; Addgene and depositors for sharing plasmids; Biorender for image creation; and Mr. Greg Sabat for mass spectrometry analysis and interpretation.

## Author Contributions Statement

Biological Experiments: ARP, EGB, SM, IS, SIA, SC, HA, MG, SB, JK

Conceptualization: ARP, MPR, JK

Writing, Reviewing, Editing: ARP, MPR, JK

Funding: MPR, JK

## References

Almeida, M. P., Welker, J. M., Siddiqui, S., Luiken, J., Ekker, S. C., Clark, K. J., Essner, J. J. and McGrail, M. (2021) ‘Endogenous zebrafish proneural Cre drivers generated by CRISPR/Cas9 short homology directed targeted integration’, Sci Rep 11(1): 1732.

Aranda, R. th, Cai, H., Worley, C. E., Levin, E. J., Li, R., Olson, J. S., Phillips, G. N., Jr. and Richards, M. P. (2009) ‘Structural analysis of fish versus mammalian hemoglobins: effect of the heme pocket environment on autooxidation and hemin loss’, Proteins 75(1): 217–30.

Ata, H., Clark, K. J. and Ekker, S. C. (2016) ‘The zebrafish genome editing toolkit’, Methods Cell Biol 135: 149–70.

Auer, T. O., Duroure, K., De Cian, A., Concordet, J. P. and Del Bene, F. (2014) ‘Highly efficient CRISPR/Cas9-mediated knock-in in zebrafish by homology-independent DNA repair’, Genome Res 24(1): 142–53.

Babaei, F., Ramalingam, R., Tavendale, A., Liang, Y., Yan, L. S., Ajuh, P., Cheng, S. H. and Lam, Y. W. (2013) ‘Novel blood collection method allows plasma proteome analysis from single zebrafish’, J Proteome Res 12(4): 1580–90.

Balasubramanian, S., Rangasamy, S., Vivekanandam, R. and Perumal, E. (2023) ‘Acute exposure to tenorite nanoparticles induces phenotypic and behavior alterations in zebrafish larvae’, Chemosphere 339: 139681.

Brownlie, A., Hersey, C., Oates, A. C., Paw, B. H., Falick, A. M., Witkowska, H. E., Flint, J., Higgs, D., Jessen, J., Bahary, N. et al. (2003) ‘Characterization of embryonic globin genes of the zebrafish’, Dev Biol 255(1): 48–61.

Burg, L., Palmer, N., Kikhi, K., Miroshnik, E. S., Rueckert, H., Gaddy, E., MacPherson Cunningham, C., Mattonet, K., Lai, S. L., Marin-Juez, R. et al. (2018) ‘Conditional mutagenesis by oligonucleotide-mediated integration of loxP sites in zebrafish’, PLoS Genet 14(11): e1007754.

Candido-Ferreira, I. L., Lukoseviciute, M. and Sauka-Spengler, T. (2023) ‘Multi-layered transcriptional control of cranial neural crest development’, Semin Cell Dev Biol 138: 1–14.

Chan, F. Y., Robinson, J., Brownlie, A., Shivdasani, R. A., Donovan, A., Brugnara, C., Kim, J., Lau, B. C., Witkowska, H. E. and Zon, L. I. (1997) ‘Characterization of adult alpha- and beta-globin genes in the zebrafish’, Blood 89(2): 688–700.

Curran, K., Raible, D. W. and Lister, J. A. (2009) ‘Foxd3 controls melanophore specification in the zebrafish neural crest by regulation of Mitf’, Dev Biol 332(2): 408–17.

Engler, C., Kandzia, R. and Marillonnet, S. (2008) ‘A one pot, one step, precision cloning method with high throughput capability’, PLoS One 3(11): e3647.

Gutierrez-Triana, J. A., Tavhelidse, T., Thumberger, T., Thomas, I., Wittbrodt, B., Kellner, T., Anlas, K., Tsingos, E. and Wittbrodt, J. (2018) ’Efficient single-copy HDR by 5’ modified long dsDNA donors’, Elife 7.

Hochgreb-Hagele, T. and Bronner, M. E. (2013) ‘A novel FoxD3 gene trap line reveals neural crest precursor movement and a role for FoxD3 in their specification’, Dev Biol 374(1): 1–11.

Hoshijima, K., Jurynec, M. J., Klatt Shaw, D., Jacobi, A. M., Behlke, M. A. and Grunwald, D. J. (2019) ‘Highly Efficient CRISPR-Cas9-Based Methods for Generating Deletion Mutations and F0 Embryos that Lack Gene Function in Zebrafish’, Dev Cell 51(5): 645–657 e4.

Howe, K. Clark, M. D. Torroja, C. F. Torrance, J. Berthelot, C. Muffato, M. Collins, J. E. Humphray, S. McLaren, K. Matthews, L., et al. (2013) ‘The zebrafish reference genome sequence and its relationship to the human genome’, Nature 496(7446): 498–503.

Hwang, W. Y., Fu, Y., Reyon, D., Maeder, M. L., Tsai, S. Q., Sander, J. D., Peterson, R. T., Yeh, J. R. and Joung, J. K. (2013) ‘Efficient genome editing in zebrafish using a CRISPR-Cas system’, Nat Biotechnol 31(3): 227–9.

Kassa, T., Whalin, J. G., Richards, M. P. and Alayash, A. I. (2021) ‘Caffeic acid: an antioxidant with novel antisickling properties’, FEBS Open Bio 11(12): 3293–3303.

Kay, M. A., He, C. Y. and Chen, Z. Y. (2010) ‘A robust system for production of minicircle DNA vectors’, Nat Biotechnol 28(12): 1287–9.

Keating, M., Hagle, R., Osorio-Mendez, D., Rodriguez-Parks, A., Almutawa, S. I. and Kang, J. (2024) ‘A robust knock-in approach using a minimal promoter and a minicircle’, Dev Biol 505: 24–33.

Lee, H. C., Huang, H. Y., Lin, C. Y., Chen, Y. H. and Tsai, H. J. (2006) ‘Foxd3 mediates zebrafish myf5 expression during early somitogenesis’, Dev Biol 290(2): 359–72.

Levic, D. S., Yamaguchi, N., Wang, S., Knaut, H. and Bagnat, M. (2021) ‘Knock-in tagging in zebrafish facilitated by insertion into non-coding regions’, Development 148(19).

Lister, J. A., Cooper, C., Nguyen, K., Modrell, M., Grant, K. and Raible, D. W. (2006) ‘Zebrafish Foxd3 is required for development of a subset of neural crest derivatives’, Dev Biol 290(1): 92–104.

Liu, F., Kambakam, S., Almeida, M. P., Ming, Z., Welker, J. M., Wierson, W. A., Schultz-Rogers, L. E., Ekker, S. C., Clark, K. J., Essner, J. J. et al. (2022) ‘Cre/lox regulated conditional rescue and inactivation with zebrafish UFlip alleles generated by CRISPR-Cas9 targeted integration’, Elife 11.

Liu, Y., Li, C., Qiu, Y., Chen, S., Luo, Y., Xiong, D., Zhao, J., Ye, J., Wang, X., Qin, W. et al. (2025) ’Base editors in zebrafish: a new era for functional genomics and disease modeling’, Front Genome Ed 7: 1598887.

Manning, J. M., Manning, L. R., Dumoulin, A., Padovan, J. C. and Chait, B. (2020) ‘Embryonic and Fetal Human Hemoglobins: Structures, Oxygen Binding, and Physiological Roles’, Subcell Biochem 94: 275–296.

Mi, J. and Andersson, O. (2023) ‘Efficient knock-in method enabling lineage tracing in zebrafish’, Life Sci Alliance 6(5).

Mosimann, C., Kaufman, C. K., Li, P., Pugach, E. K., Tamplin, O. J. and Zon, L. I. (2011) ‘Ubiquitous transgene expression and Cre-based recombination driven by the ubiquitin promoter in zebrafish’, Development 138(1): 169–77.

Odenthal, J. and Nusslein-Volhard, C. (1998) ‘fork head domain genes in zebrafish’, Dev Genes Evol 208(5): 245–58.

Oikemus, S., Hu, K., Shin, M., Idrizi, F., Goodman-Khan, A., Kolb, A., Ghanta, K. S., Lee, J., Wagh, A., Wolfe, S. A. et al. (2025) ‘Identifying optimal conditions for precise knock-in of exogenous DNA into the zebrafish genome’, Development 152(12).

Petri, K., Zhang, W., Ma, J., Schmidts, A., Lee, H., Horng, J. E., Kim, D. Y., Kurt, I. C., Clement, K., Hsu, J. Y. et al. (2022) ‘CRISPR prime editing with ribonucleoprotein complexes in zebrafish and primary human cells’, Nat Biotechnol 40(2): 189–193.

Prykhozhij, S. V. and Berman, J. N. (2024) ‘Mutation Knock-in Methods Using Single-Stranded DNA and Gene Editing Tools in Zebrafish’, Methods Mol Biol 2707: 279–303.

Prykhozhij, S. V., Fuller, C., Steele, S. L., Veinotte, C. J., Razaghi, B., Robitaille, J. M., McMaster, C. R., Shlien, A., Malkin, D. and Berman, J. N. (2018) ‘Optimized knock-in of point mutations in zebrafish using CRISPR/Cas9’, Nucleic Acids Res 46(17): 9252.

Qin, W., Liang, F., Lin, S. J., Petree, C., Huang, K., Zhang, Y., Li, L., Varshney, P., Mourrain, P., Liu, Y. et al. (2024) ‘ABE-ultramax for high-efficiency biallelic adenine base editing in zebrafish’, Nat Commun 15(1): 5613.

Richardson, C., Kelsh, R. N. and R. J. Richardson (2023) ‘New advances in CRISPR/Cas-mediated precise gene-editing techniques’, Dis Model Mech 16(2).

Rosello, M., Serafini, M., Mignani, L., Finazzi, D., Giovannangeli, C., Mione, M. C., Concordet, J. P. and Del Bene, F. (2022) ’Disease modeling by efficient genome editing using a near PAM-less base editor in vivo’, Nat Commun 13(1): 3435.

Rosello, M., Vougny, J., Czarny, F., Mione, M. C., Concordet, J. P., Albadri, S. and Del Bene, F. (2021) ‘Precise base editing for the in vivo study of developmental signaling and human pathologies in zebrafish’, Elife 10.

Sander, J. D. and Joung, J. K. (2014) ‘CRISPR-Cas systems for editing, regulating and targeting genomes’, Nat Biotechnol 32(4): 347–55.

Sauer, B. and Henderson, N. (1988) ‘Site-specific DNA recombination in mammalian cells by the Cre recombinase of bacteriophage P1’, Proc Natl Acad Sci U S A 85(14): 5166–70.

Sharrock, A. V., Mulligan, T. S., Hall, K. R., Williams, E. M., White, D. T., Zhang, L., Emmerich, K., Matthews, F., Nimmagadda, S., Washington, S. et al. (2022) ‘NTR 2.0: a rationally engineered prodrug-converting enzyme with substantially enhanced efficacy for targeted cell ablation’, Nat Methods 19(2): 205–215.

Shin, M., Yin, H. M., Shih, Y. H., Nozaki, T., Portman, D., Toles, B., Kolb, A., Luk, K., Isogai, S., Ishida, K. et al. (2023) ‘Generation and application of endogenously floxed alleles for cell-specific knockout in zebrafish’, Dev Cell.

Stewart, R. A., Arduini, B. L., Berghmans, S., George, R. E., Kanki, J. P., Henion, P. D. and Look, A. T. (2006) ‘Zebrafish foxd3 is selectively required for neural crest specification, migration and survival’, Dev Biol 292(1): 174–88.

Sugimoto, K., Hui, S. P., Sheng, D. Z. and Kikuchi, K. (2017) ‘Dissection of zebrafish shha function using site-specific targeting with a Cre-dependent genetic switch’, Elife 6.

Suzuki, K., Tsunekawa, Y., Hernandez-Benitez, R., Wu, J., Zhu, J., Kim, E. J., Hatanaka, F., Yamamoto, M., Araoka, T., Li, Z. et al. (2016) ‘In vivo genome editing via CRISPR/Cas9 mediated homology-independent targeted integration’, Nature 540(7631): 144–149.

Tessadori, F., Roessler, H. I., Savelberg, S. M. C., Chocron, S., Kamel, S. M., Duran, K. J., van Haelst, M. M., van Haaften, G. and Bakkers, J. (2018) ‘Effective CRISPR/Cas9-based nucleotide editing in zebrafish to model human genetic cardiovascular disorders’, Dis Model Mech 11(10).

Varshney, G. K., Pei, W., LaFave, M. C., Idol, J., Xu, L., Gallardo, V., Carrington, B., Bishop, K., Jones, M., Li, M. et al. (2015) ‘High-throughput gene targeting and phenotyping in zebrafish using CRISPR/Cas9’, Genome Res 25(7): 1030–42.

Wierson, W. A., Welker, J. M., Almeida, M. P., Mann, C. M., Webster, D. A., Torrie, M. E., Weiss, T. J., Kambakam, S., Vollbrecht, M. K., Lan, M. et al. (2020) ‘Efficient targeted integration directed by short homology in zebrafish and mammalian cells’, Elife 9.

Wu, Q. and Bazzini, A. A. (2023) ‘Translation and mRNA Stability Control’, Annu Rev Biochem 92: 227–245.

Yin, L., Maddison, L. A., Li, M., Kara, N., LaFave, M. C., Varshney, G. K., Burgess, S. M., Patton, J. G. and Chen, W. (2015) ‘Multiplex Conditional Mutagenesis Using Transgenic Expression of Cas9 and sgRNAs’, Genetics 200(2): 431–41.

Zhang, Y., Marshall-Phelps, K. and de Almeida, R. G. (2023) ‘Fast, precise and cloning-free knock-in of reporter sequences in vivo with high efficiency’, Development 150(12).

Zheng, S., Liang, F., Zhang, Y., Fei, J. F., Qin, W. and Liu, Y. (2023) ‘Efficient PAM-Less Base Editing for Zebrafish Modeling of Human Genetic Disease with zSpRY-ABE8e’, J Vis Exp(192).

Zhong, Z., Hu, X., Zhang, R., Liu, X., Chen, W., Zhang, S., Sun, J. and Zhong, T. P. (2025) ’Improving precision base editing of the zebrafish genome by Rad51DBD-incorporated single-base editors’, J Genet Genomics 52(1): 105–115.

