## Supplementary information for "Advancing Knock-In Approaches for Robust Genome Editing in Zebrafish"

### Supplementary Data

### Supplementary Figures and Figure Legends

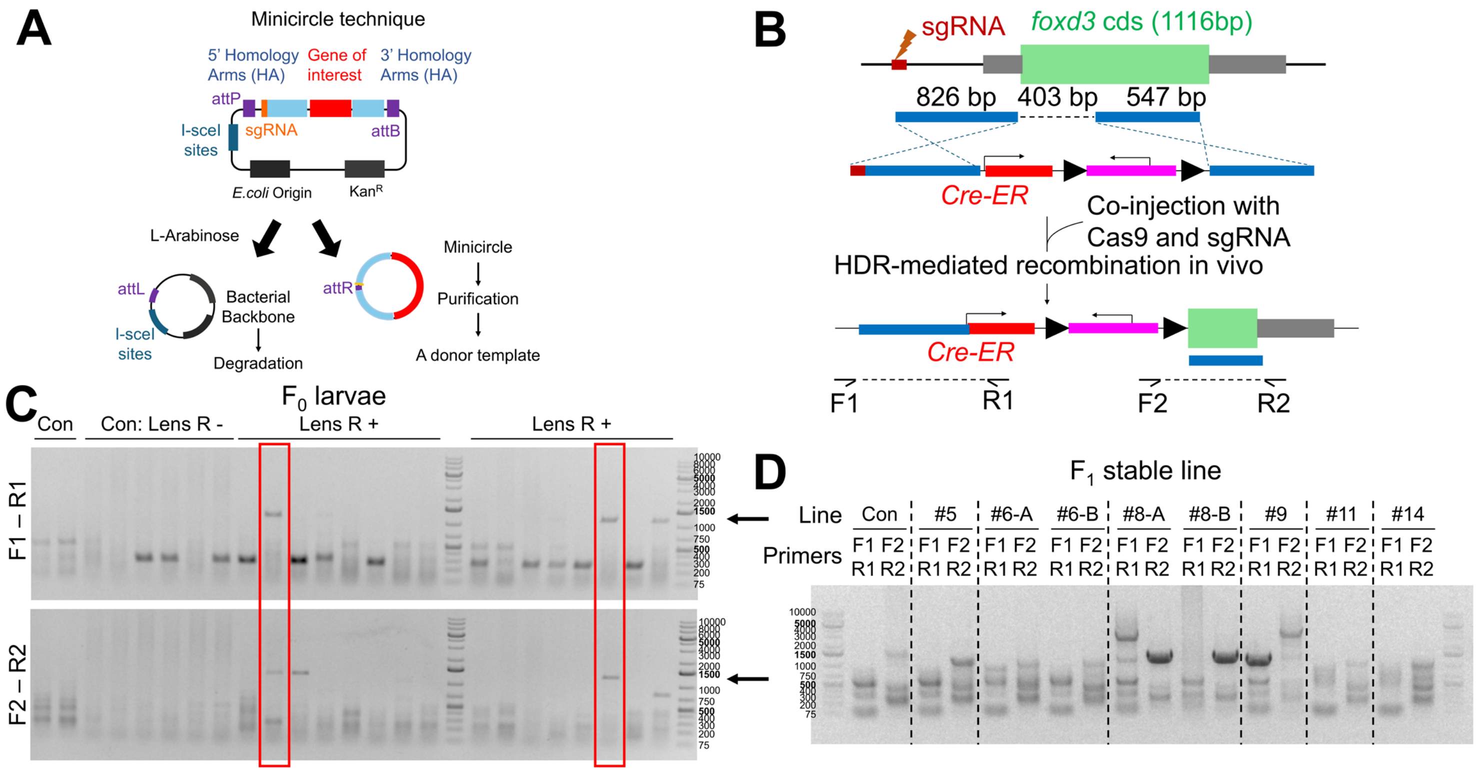


**Supplementary Fig. S1. *foxd3 CreER* knock-in reporter generation.** (**A**) Schematic of minicircle generation (**B**) Schematic of genome editing strategy to create *foxd3*$^{CreER}$ KI line. (**C**) PCR genotyping of $F_0$ larvae targeting the upstream (F1–R1) and downstream (F2–R2) regions flanking the integration site at the *foxd3* locus. (**D**) PCR genotyping of $F_1$ stable line targeting the upstream (F1–R1) and downstream (F2–R2) regions flanking the integration site at the *foxd3* locus.

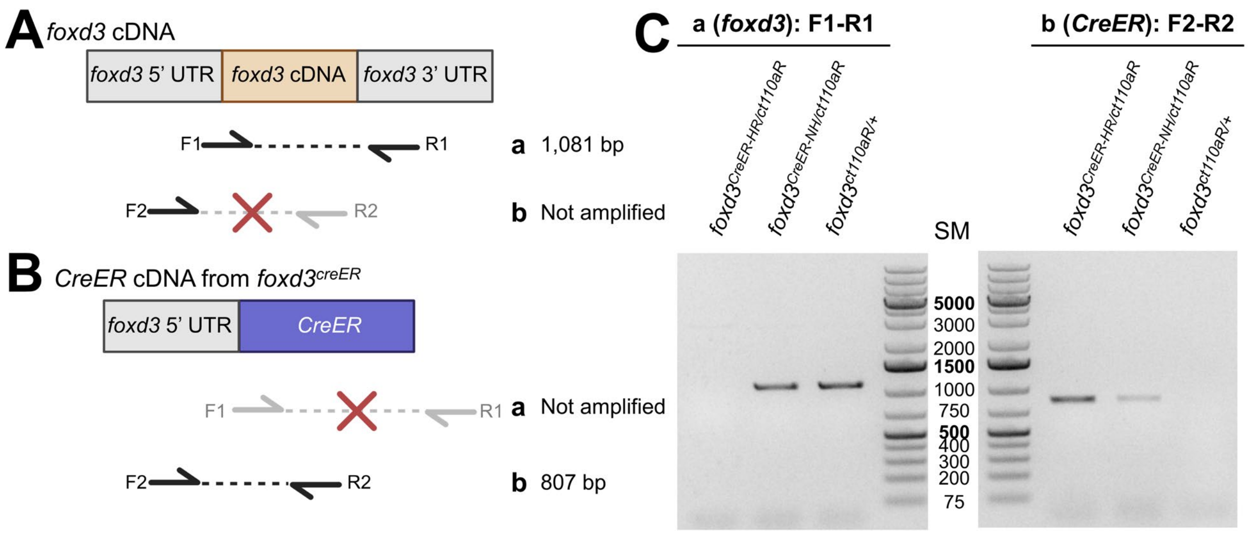


**Supplementary Fig. S2. *foxd3*$^{\textit{CreER-HR}}$ lacks *foxd3* expression.** (**A, B**) cDNA structure of *foxd3* (**A**) and *CreER* of *foxd3*$^{\textit{CreER}}$ KI lines (**B**). (**C**) RT-PCR analysis indicates lack of *foxd3* cDNA expression in *foxd3*$^{\textit{CreER-HR}}$, but not *foxd3*$^{\textit{CreER-NH}}$ (primer set #a). *CreER* cDNA is detectable in both *foxd3*$^{\textit{CreER-NH}}$ and *foxd3*$^{\textit{CreER-HR}}$ lines (primer set #b).

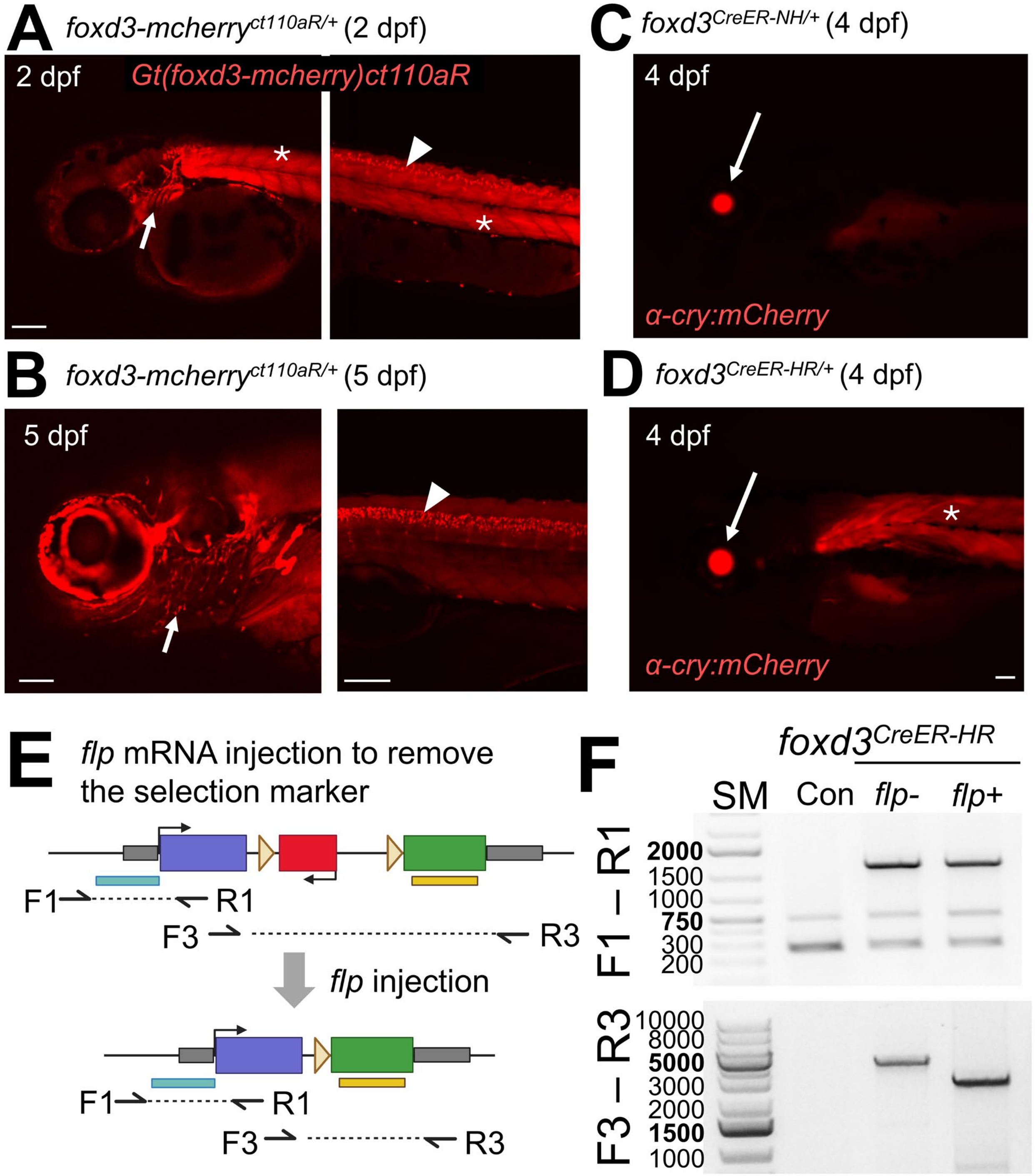


**Supplementary Fig. S3. Expression pattern of the *α-cry:mCherry* selection marker in *foxd3* CreER KI lines** (**A, B**) Expression pattern of G*t(foxd3:mcherry)$^{ct110aR}$*, a *foxd3* enhancer trap line, at 2 and 5 dpf. mCherry is detectable in NCCs, including craniofacial NCCs (arrows) and peripheral glial cells (arrowheads), as well as paraxial mesoderm (asterisks). Expression in the paraxial mesoderm declines by 5 dpf. (**C, D**) Expression pattern of the *α-cry:mCherry* selection marker in *foxd3*$^{CreER\text{-}NH/+}$ (**C**) and *foxd3*$^{CreER\text{-}HR/+}$ (**D**). While lens expression in observed in both lines (arrows), additional paraxial mesoderm expression is found in *foxd3*$^{CreER\text{-}HR/+}$ (asterisk). (**E**) Schematic of removing *FRT*-franked *α-cry:mCherry* selection marker cassette by *flp* mRNA injection (**F**) PCR genotyping of *flp*-injected larvae confirmed the deletion of the selection marker. Primer target sites are shown in **E**.

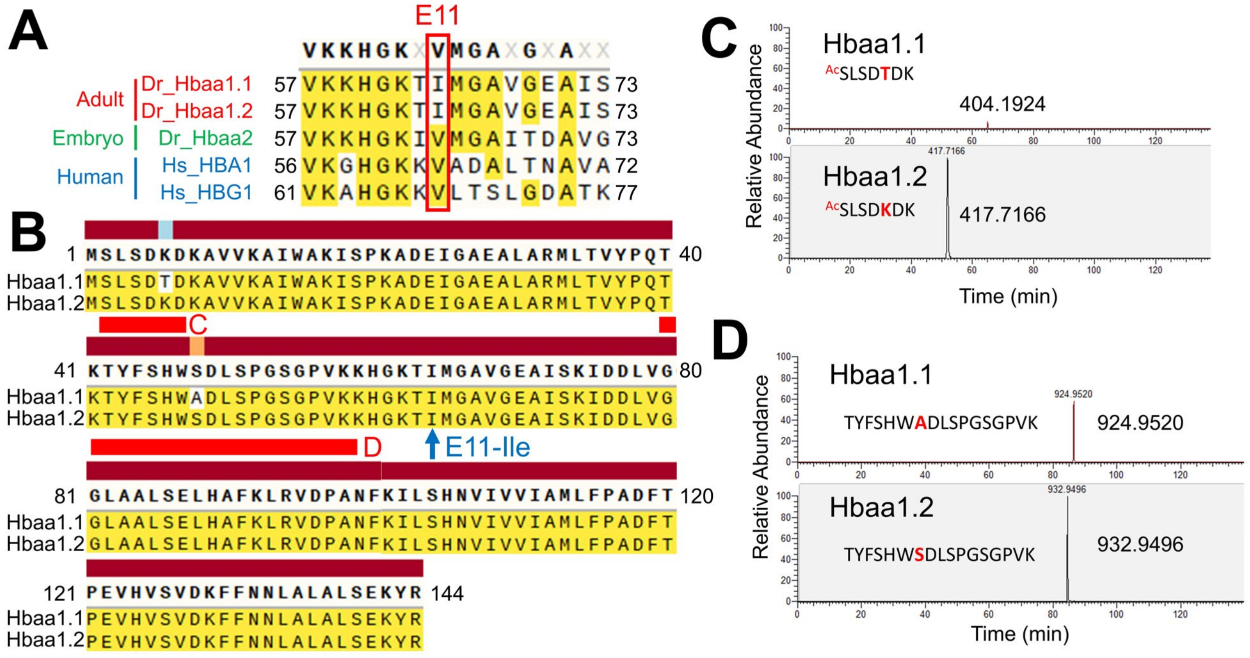


**Supplementary Fig. S4. Adult hemoglobin alpha genes in the zebrafish genome.** (**A**) amino acid alignments near the targeted E11 region of adult and embryonic hemoglobin genes in zebrafish and humans. **(B)** a.a. sequence alignment between Hbaa1.1 and Hbaa1.2 reveals two a.a. differences. (**C, D**) Mass spectrometry analysis quantifying peptides derived from Hbaa1.1 and Hbaa1.2 in adult zebrafish blood. The unique peptide sequences used for quantification are indicated in (**B**).

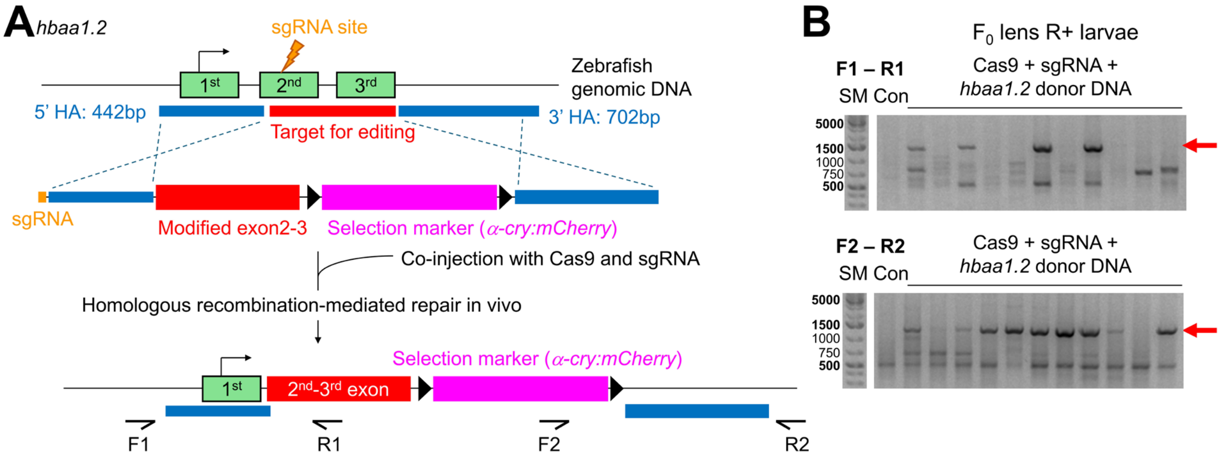


**Supplementary Fig. S5. Genome engineering to substitute I to V in Hbaa1.2.** (**A**) Schematic of genome editing strategy to create *hbaa1.2*$^{ItoV}$ allele. (**B**) PCR genotyping of $F_0$ larvae targeting the upstream (F1–R1) and downstream (F2–R2) regions flanking the integration site at the *hbaa1.2* locus.

**Supplementary Information**

**mini-golden system to assemble a donor construct for generating knock-in lines**

1. Choose your knock-in (KI) strategy
a) Targeting upstream of ATG (promoter region or 5’ untranslated region (UTR))
- Pros: Avoids concerns about out-of-frame editing.
- Con: If multiple transcription start sites exist, the resulting KI may not recapitulate expression of all isoforms.

b) Targeting within an exon or 3’ end
- Pros: Enables faithful recapitulation of endogenous expression.
- Con: Expression depends on in-frame integration; out-of-frame edits will not result in functional reporter expression.

2. Select sgRNA site
Two options are available.

a) sgRNA site located within the 5′ homology arm (HA)
- The 5′ HA typically begins at the sgRNA target site.
- After editing, the sgRNA site in the genome is often mutated, preventing re-cutting.

b) sgRNA site is located between 5’ and 3’ HAs
- The edited genome will not retain the sgRNA site, avoiding re-cleavage.
- The optimal distance between the 5′ and 3′ HAs for efficient editing remains uncertain. In our experience, HDR-mediated recombination can occur with up to a 700 bp interval between the two HAs.
- The optimal distance between cutting site and HAs remains uncertain. Based on recent study (Oikemus et al., 2025), longer than 5 bps may result in poor outcome. Thus, it is recommended to place at least one of HAs close to the cutting site.

Note: Efficient sgRNA can be predicted via CRISPRscan (https://www.crisprscan.org/) (Moreno-Mateos et al., 2015)

3. Choose middle entry vector
Select an appropriate middle entry vector based on your experimental needs:

- Vectors containing 2A sequences:
  pMC-ME plasmids with *EGFP*, *mCherry*, or *BFP* may lack the GSG linker sequence upstream of the 2A peptide. Please check the vector map.

- Vectors containing splicing acceptors (SA):
  These vectors mimic enhancer or gene trap strategies. SA sequence is derived from (Ichino et al., 2020).

4. Design primers with BsaI enzyme sites

Design primers to amplify 5′ and 3′ HAs with appropriate overhangs and BsaI recognition sites. If a BsaI site exists within your HA, it must be mutated. In-fusion PCR can be used for this purpose.

Note: The sgRNA site in the minicircle will be cleaved in the cell, generating a linearized donor construct.

*Primer structure*
5' HA CAGT forward primer: atc ggtctc t CAGT (sgRNA sequence of target region + PAM) (target region 5’ forward sequence)
5' HA TCGC reverse primer: gat GGTCTC T GCGA (target region 5’ reverse sequence)
3' HA CTCA forward primer: atc GGTCTC T CTCA (target region 3’ forward sequence)
3' HA CCAA reverse primer: gat GGTCTC T TTGG (target region 3’ reverse sequence)

5. Amplify homology arm fragments
Run PCR using primers and genomic DNA extracted from fish that will be used for injection. Purify PCR products for 5’ and 3’ HA fragments via gel extraction approach.

6. Perform Golden-Gate reaction
Set up the Golden Gate ligation reaction with the following components:
- 1 µl T4 DNA ligase buffer
- 25–50 ng each of 5′ and 3′ HA cassettes
- 100 ng chosen middle entry (pMC-ME) vector
- 100 ng destination vector
- 0.5 µl BsaI-HF v2
- 0.5 µl T4 DNA ligase
- Add nuclease-free water to a final volume of 10 µl

7. Incubate mixture 6x (37ºC for 20 min, 16ºC for 15 min), followed by 37ºC for 30 min and 80ºC for 15 min.

8. The reaction is ready for transformation (use 5 µl of the ligation and plate 20% of the transformants). Transform and spread onto kanamycin (100 µg/ml) plates

9. Screen positive colonies by colony PCR.

10. Purify a plasmid and confirm it by enzyme mapping or sequencing

11. Perform minicircle DNA purification and inject into the one-cell stage zebrafish embryos. See the details in (Keating et al., 2024)

12. Final vector sequence order

Version 1
vector - CAGT - sgRNA – 5’ HA – AGTC – pME – CTCA – 3’ HA – CCAA – vector

Version 2
vector - CAGT - sgRNA – 5’ HA – TGCG – pME – CTCA – 3’ HA – CCAA – vector

**mini-golden system to assemble a donor construct for single amino acid change**

1. Select an sgRNA near the target amino acid
Note: Efficient sgRNA can be predicted via CRISPRscan (https://www.crisprscan.org/) (Moreno-Mateos et al., 2015)

2. Validate sgRNA cutting efficiency
Inject sgRNA and Cas9 into embryos, followed by an sgRNA efficiency assay (e.g., T7 endonuclease assay) to assess cleavage at the target site.

3. Modify codons using synonymous substitutions.
Modify codons from the target amino acid (or from the sgRNA site) to the end of the same exon using synonymous codons. Codon replacement may lead to unexpected outcomes, such as affecting mRNA stability (Wu and Bazzini, 2023). Therefore, minimizing replacement is important. To address this, we follow these guidelines:

- Swapping: Replace codons with synonymous alternatives that maintain the overall codon usage balance. For example, glycine (G) can be encoded by GGT, GGC, GGA, or GGG. If the target exon contains two instances each of GGC and GGG, we may swap GGC with GGG to preserve codon composition.
- Altering: When swapping is impractical, we substitute with the most frequently used synonymous codon based on the zebrafish codon usage table (Subramanian et al., 2022).
- Unchanged: Codons encoding methionine (M) and tryptophan (W), which each have only a single codon, will remain unaltered.

4. Complete the template (**Supplementary File 13**) and design a synthetic exon that minimizes codon altering but prioritizes codon swapping where possible.

5. Choose 5' and 3' homology arm (HA) region.
Note*:* The Kang lab typically uses 400–500 bp fragments for each HA.

6. The synthetic fragment information: contain sgRNA, 5' HA, and a synthetic exon

Fragment structure:
atc ggtctct (BsaI site) - CAGT – sgRNA + PAM – 5' HA – altered target amino acid – codon-substituted region – TCGC agagacc (BsaI site) atc

7. To detect whether synthetic exon sequences generate de novo splicing donor or acceptor site, it is recommended to review the sequence using software, such as spliceator (https://www.lbgi.fr/spliceator/) (Scalzitti et al., 2021). If score is high, change to other synonymous alternatives.

8. Perform in siloco subcloning
Simulate the cloning strategy to ensure proper assembly and sequence integrity.

Vector - CAGT - sgRNA – 5’ HA – altered amino acid – a codon substituted region TCGC – pMid_acrymCherry-FRT – CTCA – 3’ HA – CCAA – vector

9. Synthesize a synthetic exon (e.g. IDT gBlock option) and order primers from the company

*Gene fragment for synthesis*
atc ggtctct (BsaI site) - CAGT - sgRNA – 5’ HA – Altered amino acid – a codon substituted region – TCGC agagacc (BsaI site) atc

*Primer structure*
5' HA CAGT forward primer: atc ggtctc t CAGT (sgRNA sequence of target region) (target region 5’ forward sequence)
5' HA TCGC reverse primer: gat GGTCTC T GCGA (target region 5’ reverse complement sequence)
3' HA CTCA forward primer: atc GGTCTC T CTCA (target region 3’ forward sequence)
3' HA CCAA reverse primer: gat GGTCTC T TTGG (target region 3’ reverse complement sequence)

10. Perform PCR to get 5’HA-synthetic-exon and 3’ HA, and assemble all fragments into pDest vector via Golden gate reaction

- 1 µl T4 DNA ligase buffer
- 25–50 ng each of 5′ HA-synthetic exon fragment and 3′ HA cassette
- 100 ng pMC-ME-v2_FRT-acrymCherry-FRT
- 100 ng destination vector
- 0.5 µl BsaI-HF v2
- 0.5 µl T4 DNA ligase
- Add nuclease-free water to a final volume of 10 µl

Note: 5’ HA-synthetic exon can be obtained by PCR using synthetized fragment as a template
Note: 3’ HA can be obtained by PCR using gDNA as a template

Incubate mixture 8x (37ºC for 20 min, 16ºC for 15 min), followed by 37ºC for 30 min and 80ºC for 15 min.

11. The reaction is ready for transformation (use 5 µl of the ligation and plate 20% of the transformants). Transform and spread onto kanamycin (100 µg/ml) plates

12. Screen positive colonies by colony PCR.

13. Purify a plasmid and confirm it by enzyme mapping or sequencing

14. Perform minicircle DNA purification and inject into the one-cell stage zebrafish embryos. See the details in (Keating et al., 2024).

**Supplementary Table S1. Plasmid list of the mini-golden**

| Plasmid Name | Addgene Plasmid ID | Linker between BsaI |
|---|---|---|
| pMC-Dest-BsaI | 200550 | CAGT-CCAA |
| pMC-ME-v1_EGFP-pA | 200513 | AGTC-CTCA |
| pMC-ME-v1_2a-EGFP-pA ORF-1 with ATG | 200514 | AGTC-CTCA |
| pMC-ME-v1_2a-EGFP-pA ORF-2 with ATG | 200515 | AGTC-CTCA |
| pMC-ME-v1_2a-EGFP-pA ORF-3 with ATG | 200516 | AGTC-CTCA |
| pMC-ME-v1_2a-EGFP-pA ORF-1 without ATG | 200517 | AGTC-CTCA |
| pMC-ME-v1_2a-EGFP-pA ORF-2 without ATG | 200518 | AGTC-CTCA |
| pMC-ME-v1_2a-EGFP-pA ORF-3 without ATG | 200519 | AGTC-CTCA |
| pMC-ME-v1_memb-EGFP-pA | 200520 | AGTC-CTCA |
| pMC-ME-v1_mCherry-pA | 200521 | AGTC-CTCA |
| pMC-ME-v1_2a-mCherry-pA ORF-1 with ATG | 200522 | AGTC-CTCA |
| pMC-ME-v1_2a-mCherry-pA ORF-2 with ATG | 200523 | AGTC-CTCA |
| pMC-ME-v1_2a-mCherry-pA ORF-3 with ATG | 200524 | AGTC-CTCA |
| pMC-ME-v1_2a-mCherry-pA ORF-1 without ATG | 200525 | AGTC-CTCA |
| pMC-ME-v1_2a-mCherry-pA ORF-2 without ATG | 200526 | AGTC-CTCA |
| pMC-ME-v1_2a-mCherry-pA ORF-3 without ATG | 200527 | AGTC-CTCA |
| pMC-ME-v1_memb-mCherry-pA | 200528 | AGTC-CTCA |
| pMC-ME-v1_mCherry-P2A-NTRv2-pA | 200529 | AGTC-CTCA |
| pMC-ME-v1_T2A-mCherry-P2A-NTRv2-pA ORF-1 | 200530 | AGTC-CTCA |
| pMC-ME-v1_T2A-mCherry-P2A-NTRv2-pA ORF-2 | 200531 | AGTC-CTCA |
| pMC-ME-v1_T2A-mCherry-P2A-NTRv2-pA ORF-3 | 200532 | AGTC-CTCA |
| pMC-ME-v1_BFP-pA | 200533 | AGTC-CTCA |
| pMC-ME-v1_2a-BFP-pA ORF-1 without ATG | 200534 | AGTC-CTCA |
| pMC-ME-v1_2a-BFP-pA ORF-2 without ATG | 200535 | AGTC-CTCA |
| pMC-ME-v1_2a-BFP-pA ORF-3 without ATG | 200536 | AGTC-CTCA |
| pMC-ME-v1_memb-BFP-pA | 200537 | AGTC-CTCA |
| pMC-ME-v1_BFP-P2A-NTRv2-pA | 200538 | AGTC-CTCA |
| pMC-ME-v1_T2A-BFP-P2A-NTRv2-pA ORF-1 | 200539 | AGTC-CTCA |
| pMC-ME-v1_T2A-BFP-P2A-NTRv2-pA ORF-2 | 200540 | AGTC-CTCA |
| pMC-ME-v1_T2A-BFP-P2A-NTRv2-pA ORF-3 | 200541 | AGTC-CTCA |
| pMC-ME-v1_CreER-pA | 200542 | AGTC-CTCA |
| pMC-ME-v1_CreER-pA-FRT-acry:venus-pA-FRT | 200543 | AGTC-CTCA |
| pMC-ME-v1_CreER-pA-FRT-acry:mCherry-pA-FRT | 200544 | AGTC-CTCA |
| pMC-ME-v1_P2A-CreER-pA ORF1 FRT-acry:mCherry-pA-FRT | 200545 | AGTC-CTCA |
| pMC-ME-v1_P2A-CreER-pA ORF2 FRT-acry:mCherry-pA-FRT | 200546 | AGTC-CTCA |

| | | |
|---|---|---|
| pMC-ME-v1_P2A-CreER-pA ORF3 FRT-acry:mCherry-pA-FRT | 200547 | AGTC-CTCA |
| pMC-Dest-uni-gRNA5-BsaI | 241150 | CAGT-CCAA |
| pMC-ME-v2_EGFP-pA | 241151 | TCGC-CTCA |
| pMC-ME-v2_memb-EGFP-pA | 241152 | TCGC-CTCA |
| pMC-ME-v2_P2A-EGFP-pA-ORF-1withoutATG | 241153 | TCGC-CTCA |
| pMC-ME-v2_P2A-EGFP-pA-ORF-2withoutATG | 241154 | TCGC-CTCA |
| pMC-ME-v2_P2A-EGFP-pA-ORF-3withoutATG | 241155 | TCGC-CTCA |
| pMC-ME-v2_mCherry-pA | 241156 | TCGC-CTCA |
| pMC-ME-v2_memb-mCherry-pA | 241157 | TCGC-CTCA |
| pMC-ME-v2_P2A-mCherry-pA-ORF-1withoutATG | 241158 | TCGC-CTCA |
| pMC-ME-v2_P2A-mCherry-pA-ORF-2withoutATG | 241159 | TCGC-CTCA |
| pMC-ME-v2_P2A-mCherry-pA-ORF-3withoutATG | 241160 | TCGC-CTCA |
| pMC-ME-v2_memb-BFP-pA | 241161 | TCGC-CTCA |
| pMC-ME-v2_pMC-ME_2a-BFP-ORF-1 withoutATG | 241162 | TCGC-CTCA |
| pMC-ME-v2_P2A-BFP-pA-ORF-2withoutATG | 241163 | TCGC-CTCA |
| pMC-ME-v2_P2A-BFP-pA-ORF-3withoutATG | 241164 | TCGC-CTCA |
| pMC-ME-v2_CreER-pA_FRT-acrymCherry | 241165 | TCGC-CTCA |
| pMC-ME-v2_CreER-pA_FRT-acry-venus | 241166 | TCGC-CTCA |
| pMC-ME-v2_P2A-CreER-pA-ORF1-FRT-acrymCherry | 241167 | TCGC-CTCA |
| pMC-ME-v2_P2A-CreER-pA-ORF2-FRT-acrymCherry | 241168 | TCGC-CTCA |
| pMC-ME-v2_P2A-CreER-pA-ORF3-FRT-acrymCherry | 241169 | TCGC-CTCA |
| pMC-ME-v2_MCS_BsmBI | 241170 | TCGC-CTCA |
| pMC-ME-v2_mNeongreen | 241171 | TCGC-CTCA |
| pMC-ME-v2_linker-P2A-lifeact-mNeongreen-ORF-1 | 241172 | TCGC-CTCA |
| pMC-ME-v2_linker-P2A-lifeact-mNeongreen-ORF-2 | 241173 | TCGC-CTCA |
| pMC-ME-v2_linker-P2A-lifeact-mNeongreen-ORF-3 | 241174 | TCGC-CTCA |
| pMC-ME-v2_memb-mNeongreen | 241175 | TCGC-CTCA |
| pMC-ME-v2_linker-P2A-mNeongreen-ORF1 | 241176 | TCGC-CTCA |
| pMC-ME-v2_linker-P2A-mNeongreen-ORF2 | 241177 | TCGC-CTCA |
| pMC-ME-v2_linker-P2A-mNeongreen-ORF3 | 241178 | TCGC-CTCA |
| pMC-ME-v2_FRT-acrymCherry-FRT | 241179 | TCGC-CTCA |
| pMC-ME-v2_linker-T2A-NTRv2-linker-P2A-mNeonGreen ORF1 | 241180 | TCGC-CTCA |
| pMC-ME-v2_linker-T2A-NTRv2-linker-P2A-mNeonGreen ORF2 | 241181 | TCGC-CTCA |
| pMC-ME-v2_linker-T2A-NTRv2-linker-P2A-mNeonGreen ORF3 | 241182 | TCGC-CTCA |
| pMC-ME-v2_linker-P2A-mem-mScarlet-ORF-1 | 241183 | TCGC-CTCA |
| pMC-ME-v2_linker-P2A-mem-mScarlet-ORF-2 | 241184 | TCGC-CTCA |
| pMC-ME-v2_linker-P2A-mem-mScarlet-ORF-3 | 241185 | TCGC-CTCA |
| pMC-ME-v2_SA-linker-P2A-mem-mScarlet-ORF-1 | 241186 | TCGC-CTCA |
| pMC-ME-v2_SA-linker-P2A-mem-mScarlet-ORF-2 | 241187 | TCGC-CTCA |

| | | |
|---|---|---|
| pMC-ME-v2_SA-linker-P2A-mem-mScarlet-ORF-3 | 241188 | TCGC-CTCA |
| pMC-ME-v2_nzCas9n-GSG-linker-P2A-mNeongreen | 241189 | TCGC-CTCA |
| pMC-ME-v2_3x-FLAG-nzdCas9-V5-APEX2n-GSG-linker-P2A-mNeongreen | 241190 | TCGC-CTCA |
| pMC-ME-v2_nAPEX2-V5-zdCas9n-3x-FLAG-GSG-linker-P2A-mNeongreen | 241191 | TCGC-CTCA |
| pMC-ME-v2_linker-T2A-NTRv2-linker-P2A-mNeonGreen-CAAXORF1 | 241192 | TCGC-CTCA |
| pMC-ME-v2_linker-T2A-NTRv2-linker-P2A-mNeonGreen-CAAXORF2 | 241193 | TCGC-CTCA |
| pMC-ME-v2_linker-T2A-NTRv2-linker-P2A-mNeonGreen-CAAXORF3 | 241194 | TCGC-CTCA |
| pMC-ME-v2_mCherry-P2A-NTRv2-pA | 241195 | TCGC-CTCA |
| pMC-ME-v2_T2A-mCherry-P2A-NTRv2-pA-ORF-1 | 241196 | TCGC-CTCA |
| pMC-ME-v2_T2A-mCherry-P2A-NTRv2-pA-ORF-2 | 241197 | TCGC-CTCA |
| pMC-ME-v2_T2A-mCherry-P2A-NTRv2-pA-ORF-3 | 241198 | TCGC-CTCA |
| pMC-ME-v2_T2A-BFP-P2A-NTRv2-pA-ORF-1 | 241199 | TCGC-CTCA |
| pMC-ME-v2_T2A-BFP-P2A-NTRv2-pA-ORF-2 | 241200 | TCGC-CTCA |
| pMC-ME-v2_T2A-BFP-P2A-NTRv2-pA-ORF-3 | 241201 | TCGC-CTCA |
| pMC-ME-v2_lifeact-mNeongreen-pA | 241202 | TCGC-CTCA |
| pMC-ME-v2_SA-TCGC-mNeongreen | 241203 | TCGC-CTCA |
| pMC-ME-v2_SA-linker-P2A-lifeact-mNeongreen-ORF-1 | 241204 | TCGC-CTCA |
| pMC-ME-v2_SA-linker-P2A-lifeact-mNeongreen-ORF-2 | 241205 | TCGC-CTCA |
| pMC-ME-v2_SA-linker-P2A-lifeact-mNeongreen-ORF-3 | 241206 | TCGC-CTCA |
| pMC-ME-v2_SA-memb-mNeongreen | 241207 | TCGC-CTCA |
| pMC-ME-v2_SA-linker-P2A-mNeongreen-ORF1 | 241208 | TCGC-CTCA |
| pMC-ME-v2_SA-linker-P2A-mNeongreen-ORF2 | 241209 | TCGC-CTCA |
| pMC-ME-v2_SA-linker-P2A-mNeongreen-ORF3 | 241210 | TCGC-CTCA |
| pMC-ME-v2_SA-EGFP-pA | 241211 | TCGC-CTCA |
| pMC-ME-v2_SA-mCherry-pA | 241212 | TCGC-CTCA |
| pMC-dest_BsaI-cmlc2:mCherry | 241213 | CAGT-CCAA |
| pMC-dest_BsaI-MCS-cloningvector forgoldengate | 241214 | CAGT-CCAA |

### Supplementary files

**Supplementary file 1. Primer sequence list**
**Supplementary file 2. *foxd3* gDNA sequence**
**Supplementary file 3. *foxd3 CreER* donor vector sequence**
**Supplementary file 4. *foxd3$^{CreER}$* line 9 sequence**
**Supplementary file 5. *foxd3$^{CreER}$* line 23 sequence**
**Supplementary file 6. *hbaa1.2* gDNA sequence**
**Supplementary file 7. *hbaa1.2* ItoV donor vector sequence**
**Supplementary file 8. *hbaa1.2$^{ItoV}$* gDNA sequence**
**Supplementary file 9. Tutorial file for mini-golden–mediated amino acid substitution: synthetic fragment design for *hbaa1.2* I-to-V substitution.**
**Supplementary file 10. Tutorial file for mini-golden–mediated amino acid substitution: a donor pMC vector sequence for *hbaa1.2* I-to-V substitution.**
**Supplementary file 11. pMC-Dest-BsaI sequence**
**Supplementary file 12. pMid_acrymCherry-FRT sequence**
**Supplementary file 13. Codon_usage_change_template**

### References


Ichino, N., Serres, M. R., Urban, R. M., Urban, M. D., Treichel, A. J., Schaefbauer, K. J., Tallant, L. E., Varshney, G. K., Skuster, K. J., McNulty, M. S. et al. (2020) 'Building the vertebrate codex using the gene breaking protein trap library', *Elife* 9.

Keating, M., Hagle, R., Osorio-Mendez, D., Rodriguez-Parks, A., Almutawa, S. I. and Kang, J. (2024) 'A robust knock-in approach using a minimal promoter and a minicircle', *Dev Biol* 505: 24-33.

Moreno-Mateos, M. A., Vejnar, C. E., Beaudoin, J. D., Fernandez, J. P., Mis, E. K., Khokha, M. K. and Giraldez, A. J. (2015) 'CRISPRscan: designing highly efficient sgRNAs for CRISPR-Cas9 targeting in vivo', *Nat Methods* 12(10): 982-8.

Oikemus, S., Hu, K., Shin, M., Idrizi, F., Goodman-Khan, A., Kolb, A., Ghanta, K. S., Lee, J., Wagh, A., Wolfe, S. A. et al. (2025) 'Identifying optimal conditions for precise knock-in of exogenous DNA into the zebrafish genome', *Development* 152(12).

Scalzitti, N., Kress, A., Orhand, R., Weber, T., Moulinier, L., Jeannin-Girardon, A., Collet, P., Poch, O. and Thompson, J. D. (2021) 'Spliceator: multi-species splice site prediction using convolutional neural networks', *BMC Bioinformatics* 22(1): 561.

Subramanian, K., Payne, B., Feyertag, F. and Alvarez-Ponce, D. (2022) 'The Codon Statistics Database: A Database of Codon Usage Bias', *Mol Biol Evol* 39(8).

Wu, Q. and Bazzini, A. A. (2023) 'Translation and mRNA Stability Control', *Annu Rev Biochem* 92: 227-245.
